# Spatial omics imaging of fresh-frozen tissue and routine FFPE histopathology on a single cancer needle core biopsy: freezing device and multimodal workflow

**DOI:** 10.1101/2023.02.11.528125

**Authors:** Miriam F. Rittel, Stefan Schmidt, Cleo-Aron Weis, Emrullah Birgin, Björn van Marwick, Matthias Rädle, Steffen J. Diehl, Nuh Rahbari, Alexander Marx, Carsten Hopf

**Author notes:** equal contribution (Miriam F. Rittel, Stefan Schmidt). Institute of Pathology, Heidelberg University Hospital, Heidelberg, Germany (Cleo-Aron Weis).

## Abstract

Complex molecular alterations underlying cancer pathophysiology are intensely studied with omics methods using bulk tissue extracts. For spatially resolved tissue diagnostics using needle biopsy cores, however, histopathological analysis using stained FFPE tissue and immuno-histochemistry (IHC) of few marker proteins is currently the main clinical focus. Today, spatial omics imaging using MSI or IRI are emerging diagnostic technologies for identification and classification of various cancer types. However, to conserve tissue-specific metabolomic states, fast, reliable and precise methods for preparation of fresh-frozen (FF) tissue sections are crucial. Such methods are often incompatible with clinical practice, since spatial metabolomics and routine histopathology of needle biopsies currently require two biopsies for FF and FFPE sampling, respectively. Therefore, we developed a device and corresponding laboratory and computational workflows for multimodal spatial omics analysis of fresh-frozen, longitudinally sectioned needle biopsies to accompany standard FFPE histopathology on the same biopsy core. As proof-of-concept, we analyzed surgical human liver cancer specimen by IRI and MSI with precise co-registration and, following FFPE processing, by sequential clinical pathology analysis on the same biopsy core. This workflow allowed spatial comparison between different spectral profiles and alterations in tissue histology, as well as direct comparison to histological diagnosis without the need of an extra biopsy.

**SIMPLE SUMMARY:** Routine clinical approaches for cancer diagnosis demand fast, cost-efficient, and reliable methods, and their implementation within clinical settings. Currently, histopathology is the golden standard for tissue-based clinical diagnosis. Recently, spatially resolved molecular profiling techniques like mass spectrometry imaging (MSI) or infrared spectroscopy imaging (IRI) have increasingly contributed to clinical research, e.g., by differentiation of cancer subtypes using molecular fingerprints. However, adoption of the corresponding workflows in clinical routine remains challenging, especially for fresh-frozen tissue specimen. Here, we present a novel device based on 3D-printing technology, which facilitates sample preparation of needle biopsies for correlated clinical tissue analysis. It enables combination of MSI and IRI on fresh-frozen clinical samples with histopathological examination of the same needle core after formalin-fixation and paraffin-embedding (FFPE). This device and workflow can pave the way for a more profound understanding of biomolecular processes in cancer and, thus, aid more accurate diagnosis.

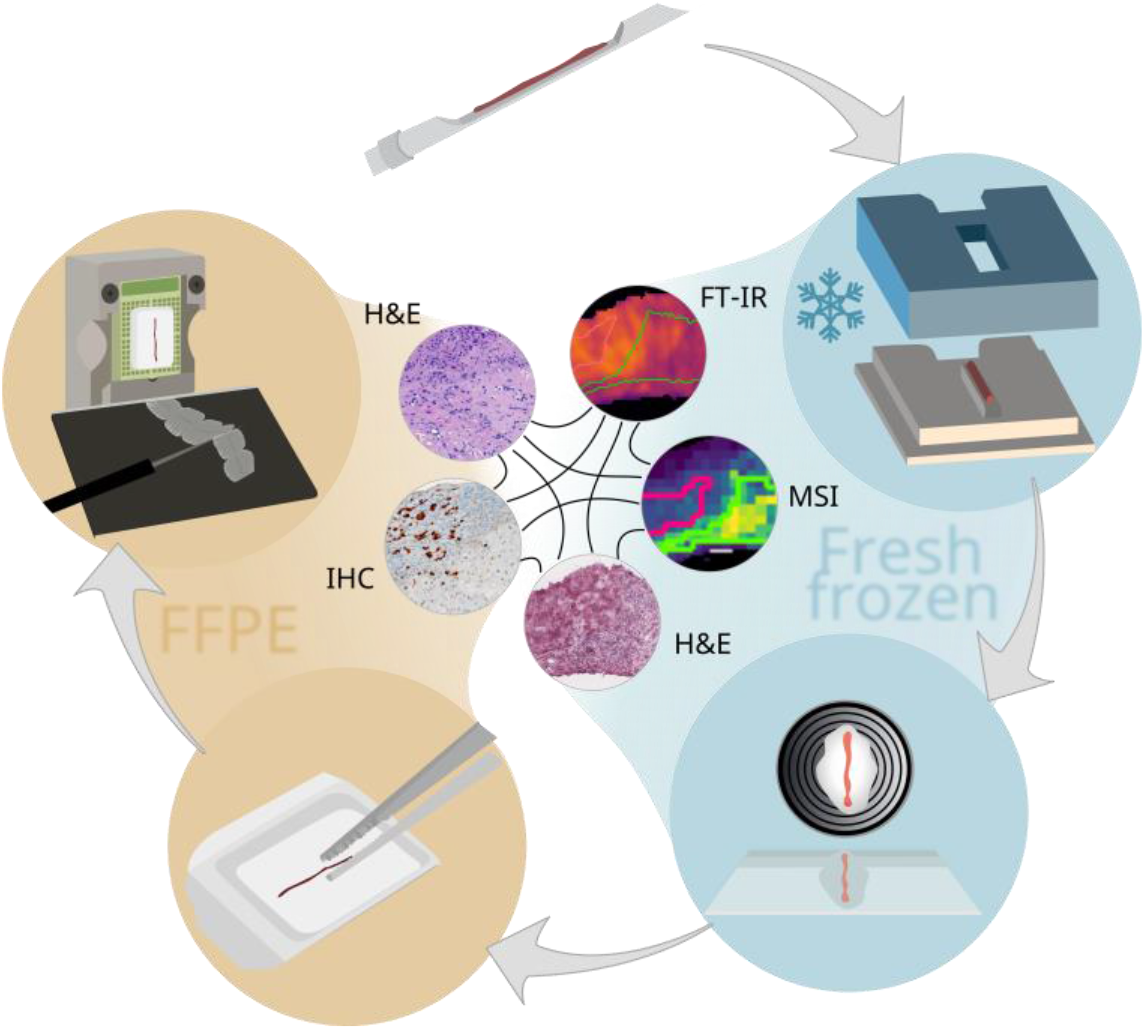

## 1. INTRODUCTION

Cancer diagnosis often requires biopsy, i.e. the extraction of small, localized tissue samples from patients using elaborated mechanical tools. Personalized therapy using interventional radiology and/or adjuvant pharmacotherapy increasingly demand advanced molecular tissue analytics, e.g. for tumor sub-classification, that can aid decision-making by tumor boards. Complementing non-invasive clinical diagnostic imaging, biopsies allow for direct investigation of the heterogeneous molecular composition and complex pathophysiology of a tissue with high cellular specificity. Core needle biopsy is a frequently used procedure for direct examination of neoplastic lesions like soft-tissue sarcoma^1^, breast cancer^2^ or liver neoplasms^3^, and for ancillary molecular testing. Pathologies of the liver such as fibrosis, cirrhosis or cancer are nowadays routinely addressed by percutaneous needle core biopsy^4^. Although this method is widely adopted in oncology and beyond, current devices are not yet designed to support- and current procedures are just starting to support modern (spatial) omics technologies^5^. In routine pre-omics era clinical practice and in accordance with clinical guidelines^6^, examination of resected tissues or biopsies is still largely performed based on expert pathology interpretation of histomorphological features of stained tissues, albeit in increasingly computational fashion^7^.. In addition, analysis of cell heterogeneity and spatial organization within the tissue can be aided by immunohistochemistry of a rather small number of marker proteins. In contrast, mass spectrometry- (MS-) based omics methodologies have recently benefitted from vastly improved sensitivity^8^, a trend that has prompted the development of miniaturized workflows with first applications for analysis of bulk extracts from core needle biopsies^9,10^.

With the advent of spatially resolved omics technologies like MSI of proteins, lipids, metabolites (“spatial lipidomics/metabolomics”)^11–14^, MSI-guided micro-proteomics^15^, microscopy-guided micro-proteomics^16^ and increasingly spatial transcriptomics^17^ and spatial multi-omics^18^ for studies of complex molecular alterations in pathophysiology in tissue specimen, the promises for future clinical diagnosis, prognosis or targeted therapy are manifold. They allow to uncover intra-tumor heterogeneity and to investigate pathophysiological processes like tumor genesis, -progression or – metastasis. In all of this, the spatial information is essential. Within the field of spatial omics, MSI provides high speed and high chemical specificity at cellular spatial resolution^11–13^. Matrix-Assisted Laser Desorption/Ionization (MALDI) MSI, in particular, has been demonstrated to be clinically relevant in multiple aspects of liver disease ranging from colorectal cancer metastasis^19^, drug distribution in liver metastases^20^ and hepatocellular carcinoma^21^ to a generally new promising technology in next-generation clinical pathology practice^22^. Moreover, studies often combine complementary imaging techniques to access the complex biomolecular context of a sample and to allow more sophisticated so-called multimodal workflows^23–26^: Label-free imaging techniques that can be performed on the same tissue section prior to MSI are infrared imaging (IRI) and IR microscopy. Due to recent technological innovations in vibrational spectroscopy, IR methods enable rapid screening and examination of tissue section for tumor margin detection, classification and grading^24,27–30^. Similar to correlative histology-based MSI approaches^31^, morphological features obtained with IR spectroscopy yield accurate labels for precise region-specific MSI-based molecular fingerprinting^24^ (“IRI-guided MSI”). However, correlating different spatial omics approaches requires sophisticated data-analysis pipelines and strategies in the field of sample preparation, image fusion, and image co-registration and data evaluation. Similar requirements are mandatory for a direct comparison with the FFPE-based histopathology, which is the current golden standard.

Whereas clinical routine analysis and (spatial) proteomics can be performed on formalin-fixed paraffin-embedded (FFPE) tissue sections, lipidomic and metabolomic approaches, including IRI and MSI, require fresh-frozen (FF) material to reveal unmodified molecular signatures. In addition, to conserve tissue-specific metabolomic states, fast, reliable and precise methods for preparation of fresh-frozen tissue sections are crucial. They are often challenging within clinical practice, however, especially for longitudinally cut biopsy sections to yield a sufficient cross-section for analysis. Processing of FF biopsy samples is difficult in terms of collection, storage, and cryo-sectioning. In addition, an optimized freezing protocol is mandatory to minimize the amount of tissue alterations due to freezing damage. Therefore, a standardized procedure limiting environmental and human error sources is advantageous. Available sample preparation protocols and tools^32^ were developed for FFPE-based tissue embedding and are inconsistent with requirements for lipid/metabolite MSI. In addition, common methodologies like freezing biopsies directly onto a metal chuck include sample transport in saline where the molecular state of the sample may be altered. Therefore, core biopsies are difficult to handle with this technique, especially for non-continuous cores or for ambitious multimodal approaches. In a classic study, Cazares and co-workers have demonstrated the combined analysis of longitudinally cut, mirrored biopsy cores by MSI and histology^33^. Shiraishi et al. have recently presented a sophisticated device to divide a biopsy into two similar parts^34^. However, the capability for snap-freezing of the sample and for analyzing both FF and subsequent FFPE tissue from the same needle core are lacking.

To overcome the challenges and limitations for use of core needle biopsies in the field of spatial omics, a device for biopsy FF sample preparation of longitudinally sectioned biopsies was developed. Clinical requirements as sterilizability, cost and easy-of-handling were also considered in the design. It allows a reliable and robust multimodal workflow to combine MALDI imaging of FF tissue with routine histopathological FFPE tissue analysis on a single biopsy core. Further, it enables handling of minimal invasive, non-continuous biopsy cores for any analyses requiring FF sample preparation. Multiple sections could be retrieved for different analyses and accurate image co-registration was employed, which allowed direct comparison. Using the presented workflow, the correlation between differentially expressed spectral profiles and the tissue histology on FF material was investigated. Furthermore, a comparison between pathological examination on FFPE and molecular or spectral profiles was performed. This study presents the design of the device and the 3D-printing manufacturing process. The corresponding workflow is described in detail. Furthermore, a feasibility study with two patients (diagnosis: hepatocellular carcinoma (HCC) and mamma metastasis in liver) and two spectral measurement techniques (MSI and Fourier transform (FT)-IRI) on FF tissue as well as clinical routine histopathology assessment on a subsequent FFPE preparation all conducted on the same biopsy core, is presented.

## 2. MATERIAL & METHODS

### 2.1. Materials

1,5-diamino-naphthalene (DAN) (D21200, Sigma-Aldrich, Taufkirchen, Germany), hydroxypropyl methylcellulose: polyvinylpyrrolidone (HPMC:PVP, home-made^35^), indium tin oxide-coated microscope slides (ITO slides) (8-12 Ohm resistance, Diamond Coatings Ltd., Brierley Hill, United Kingdom). De-ionized water (dH_2_O) was prepared in-house.

### 2.2. Tissue specimen and needle biopsies

Human tissue specimen were collected from excised tumors during surgery. Human sampling was covered by ethic vote (2012-293N-MA, 20.062012) of the Medical Ethics Committee II of the Medical Faculty Mannheim of Heidelberg University with informed consent of all participants. Following the published guidelines on the use of liver biopsy in clinical practice^6^, biopsies were excised with a core biopsy instrument (gauge size of 18G, penetration depth of 22 mm; Bard Peripheral Vascular Inc., Tempe, USA). Specimen in this study included one HCC (female, 73 years of age) and one mamma metastasis residing in liver (female, 31 years of age). Three biopsies each were taken for the HCC samples and two for the mamma metastasis sample due to size limitations of the nodule. One biopsy per entity is shown as representative example in the main manuscript; all others are depicted in the supplementary information. Initial experiments were conducted on biopsies punched from a bovine liver obtained from a local butcher.

### 2.3. Biopsy freezing device, preparation of fresh-frozen samples

Autodesk Inventor Professional 2022 (Autodesk Inc., USA) was used to compile the CAD designs of all parts required. The base compartment made from metal was directly printed as a positive module using austenitic steel powder (MetcoAdd 316L-A, Oerlikon Metco Europe GmbH, Raunheim, Switzerland) and a 3D metal printer (TruPrint 2000, Trumpf, Stuttgart, Germany). Silicone parts comprising a base- and a top compartment were retrieved from negative casting molds (**Figure 1b**). Negative casting molds were fabricated from a biocompatible photopolymer resin (FLDGOR01, Formlabs, Berlin, Germany) by low force stereolithography using a 3D-printer (Form2, Formlabs, Berlin, Germany). Prior to use, negative casting molds were washed with ethanol in an ultrasonic cleaner and subsequently dried at room temperature and ambient pressure. The two-component silicone-rubber (REPLISIL 32 N, Silconic, Lonsee, Germany) used for base was in accordance with manufacturer recommendations (solution A : solution B, 1:1 (v:v)) and poured into the casting mold directly after mixing. Forms vulcanized at room temperature (RT) and silicone curation took about 20 min. In order to use the tool for FF sample preparation, punched needle biopsies were skimmed off onto the bridge of the base compartment, which was then covered with the top compartment and filled with embedding medium (HPMC:PVP, 1:1, w/w)^35^. In brief, embedding medium was prepared by dissolving 2.5% PVP and 7.5% of HPMC in dH2O, stirred in an ice bath overnight and kept at 4 °C for several hours. For long-term storage, the hydrogel was kept at −20°C in a freezer. For snap-freezing, the biopsy-loaded device was then placed on an aluminum foil floating on liquid nitrogen. When frozen, samples were stored at −80 °C until use. For sectioning, samples were equilibrated in a cryostat (CM1860 UV, Leica Biosystems, Nussloch, Germany) at −16 °C for about 20 min. The flexible top compartment of the device was then removed, the sample was detached from the base with pre-cooled forceps and mounted upside-down onto a metal chuck for cryo-sectioning (10 μm section thickness) later used for spatial omics analyses and H&E reference staining on FF sections.

**Figure 1:**
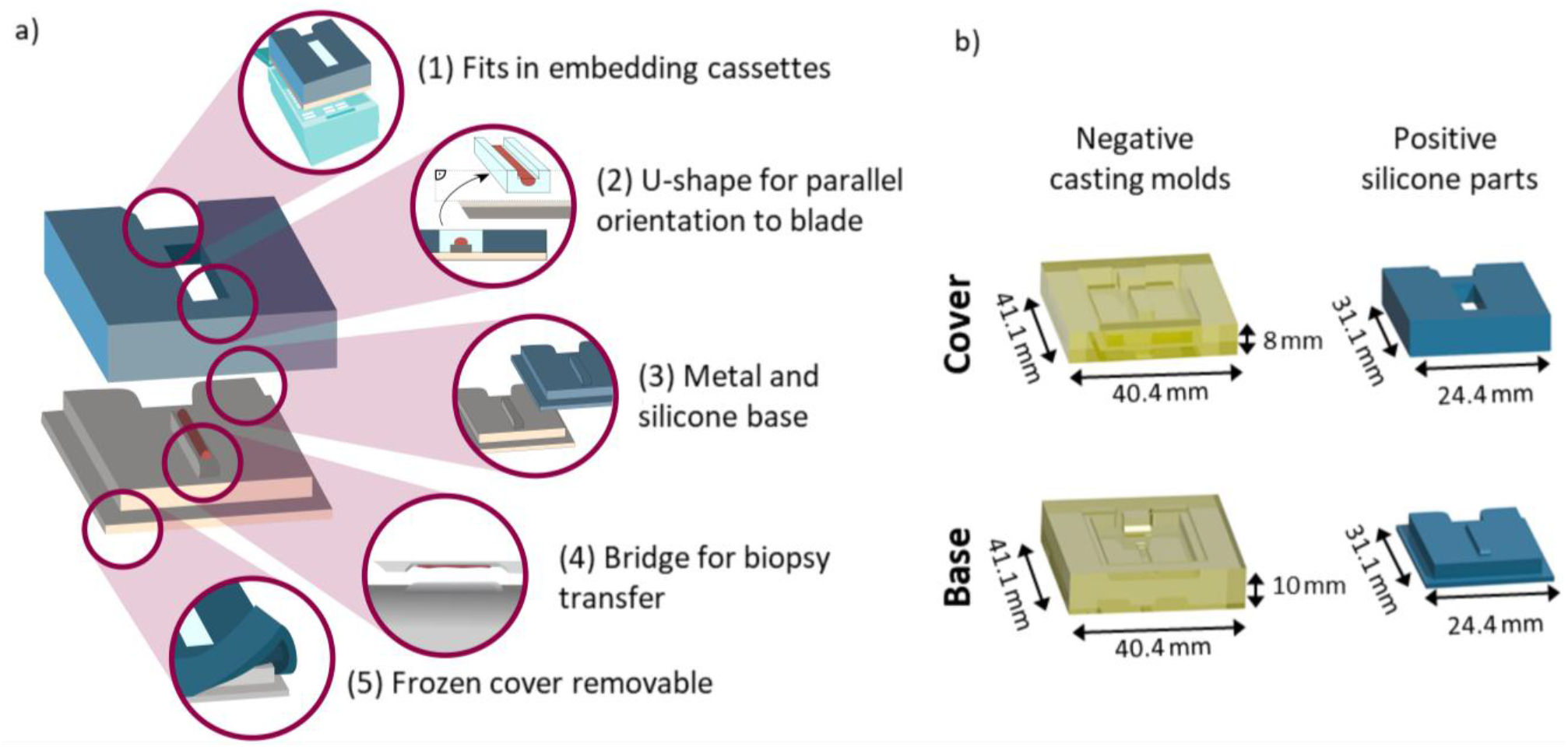
Schematic representation of features of the biopsy freezing device and conceptual production parts. **(a)** Design choices for the 3D-printed biopsy freezing device, highlighting from top to bottom: (1) Overall sizing fits into commercially available embedding cassettes for prior labelling (with pencil, aerospace markers or barcodes) of the sample. (2) U-shaped mold for precise orientation of parallel sectioning plane to blade. (3) Different base materials with different thermal conductive properties tested. (4) Bridge facilitating biopsy transfer from the needle to the device. (5) Silicone cover that is bendable at negative temperatures and therefore removable when frozen. **(b)** Technical drawings of the individually constructed parts of the device for core needle biopsy freezing including 3D-printed negative casting molds made from resin (yellow) for the cover as well as the base component, and the resulting positive silicone parts (blue) obtained from the negative molds, respectively.

### 2.4. MALDI mass spectrometry imaging (MSI) and FT-IR imaging (IRI)

For MSI and IRI, tissue cryosections were thaw-mounted onto ITO slides. Following 15 min equilibration at RT, bright-field (BF) optical reference images were acquired using an Aperio CS2 scanner (objective: 20x/0.75 NA Plan Apo, Leica Biosystems, Nussloch, Germany) for teaching of tissue positions in MSI. Subsequently, IRI was performed on a Spotlight 400 FT-IR Imaging System (Perkin Elmer LAS, Rodgau, Germany) as described elsewhere^24^. In brief, images were recorded with a spatial resolution of 25 μm, a wave number range of 750 – 4000 cm^-1^ at 4 cm^-1^ spectral resolution, 2 scans per pixel and 2.2 cm/s mirror speed. Spectra were preprocessed to common standards (2^nd^ derivative and standard normal variate (SNV) normalization)^36^, and images were visualized using *specio* as freely available python package [http://imageio.github.io]. MSI was performed on the same section after IRI. For MSI, slides were coated with DAN matrix as described elsewhere^37^. Measurements were performed on a 7 T XR magnetic resonance (MR) mass spectrometer (solariX, Bruker Daltonics, Bremen, Germany) in negative ion mode in the range of *m/z* 200 – 2000, at a lateral step-size of 50 μm, using a 1 M transient (free induction decay time of 0.98 s). On-line calibration was performed using the ubiquitous endogenous [M-H]^-^ ion of PI 38:4 (*m/z* 885.5487).

For each mass spectrum, ions from 200 shots at 2,000 Hz were accumulated at a laser power of 25 %. Prior to analysis, the system was calibrated via the ESI source with the ESI-L low concentration tuning mix (G1969-85000, Agilent Technologies, Waldbronn, Germany). Mass spectra were visualized using SCiLS Lab 2023a Pro (Bruker Daltonics). The mass-search-window of *m/z* values was inferred from the mass resolution specific for that measurement and device based on full-width at half-maximum (FWHM; resolving power of ^~^135,000 at *m/z* 400, ^~^73,500 at *m/z* 800) of the corresponding mass peak and has been determined as described^1^. MSI results were evaluated using discriminant ROC analysis and visualized using *SCiLS Lab* software (Bruker Daltonics). Molecular annotation was performed using *Metaspace* (https://metaspace2020.eu/).

### 2.5. Precise co-registration

Spectral maps from MSI and IRI were co-registered with reference pathology H&E analysis. To develop the registration workflow and assess registration quality of longitudinally sectioned biopsy samples, bovine liver samples were used. For registration, the python packages SimpleITK^38^ and simpleElastix^39^ were used. With the latter, image registration was done by sequential rigid, affine and b-spline transformations. Advanced Mattes mutual information was used as a metric for the linear interpolator^40^. In total, about 1000 iterations were performed for each linear transformation and about twice as many iterations for non-linear transformations.

Registration methods were tested and evaluated on a bovine liver data set consisting of three needle biopsy cores and then applied to the human dataset. Testing, optimization and validating of image registration parameters was performed on subsequent H&E stained bovine liver tissue sections. For visual comparison prior to image registration, color normalization between the two H&E-stained histological images was applied by aligning the color distributions of each color channel between the two images. To this end, morphological features like blood vessels were selected as landmarks based on manual examination of the tissue, and annotated using the Aperio Image Scope software (Leica Biosystems). Two different parameters were obtained to assess registration quality: a point-to-point comparison of corresponding landmarks (distances of center-of-gravity of the annotated regions) and Sorensen-Dice similarity^41^. Briefly, the formula used to calculate the DICE similarity is (DICE = 2 x overlapping area / sum of both corresponding areas). For each biopsy, six regions-of-interest (ROIs) evenly distributed over the biopsy were selected. Both parameters were calculated as a function of the relative distance between individual tissue sections. Results are presented as box-plots, with the box marking the first and third quartiles, as well as the median. Whiskers represent the highest value within the 1.5x inter-quartile-range (IQR). Results were calculated on the one hand for all landmarks of the three biopsies (total of 3 biopsies x 6 landmarks) in one plot, and on the other hand for each biopsy individually (6 landmarks per biopsy). In both cases, a second-order polynomial function was fitted to the data to guide-the-eye. Registration results are calculated and plotted using Python. Graphics were assembled using power point (Microsoft Corporation) and Inkscape (GNU General Public License, version 3).

### 2.6. Conversion of remaining FF samples to FFPE and clinical pathology analysis

After cryosectioning, the remainder of the FF biopsies in embedding medium was immersed in 5 mL of 10% phosphate-buffered formalin (Sarstedt, Nürmbrecht, Germany) at RT. Non-dissolved embedding medium could be removed during this step when floating at the top of the solution if desired. After at least 22 h, biopsies were embedded in paraffin according to the clinical routine FFPE workflow. FFPE tissue sections (2 μm thickness) obtained with a Leica RM2245 rotary microtome (Leica Biosystems) were mounted on glass slides (Superfrost Plus, Epredia, Braunschweig, Germany). Staining details for FF and FFPE prepared sections are described in **Supplementary Methods**.

## 3. RESULTS AND DISCUSSION

Solid tumors typically require biopsies for accurate diagnosis and for optimal treatment selection for individual patients. However, so far very limited (spatial) omics research has been done on biopsy cores, although proteomics, metabolomics and MSI have become mainstays in cancer research^8,12^. This is due to obstacles widely described in literature^4,42^, like insufficient handling capabilities of FF biopsy core material, which is optimal for most omics techniques. Nevertheless, clinical research consortia like the Mannheim Molecular Intervention Environment (M^2^OLIE; http://www.m2olie.de/en/) that focus on innovative tumor therapies using molecular intervention rely on the diagnosis of core biopsy material obtained in interventional radiology for diagnosis. For this reason, here a device and corresponding laboratory and computational workflows were developed, with which biopsy cores can be prepared for FF tissue handling and for subsequent reincorporation in clinical FFPE routine. Multiple sections can be retrieved for multimodal spatial omics analysis, and the remaining core can be further processed into FFPE specimen for subsequent clinical routine histopathology analysis. The overall process presented here includes the combined spectral analyses on FF cancer tissue sections and clinical routine analyses on FFPE samples on the same biopsy core for the classification of the different tissue types and the identification of morphological features within the cancer tissue section.

### 3.1. 3D-printed, flexible embedding and freezing device for longitudinal sectioning of needle biopsies

In order to facilitate parallel clinical use of single core needle biopsies in spatial multi-omics studies that require FF tissue as well as in routine histopathology utilizing FFPE tissue, we first set out to design and manufacture a new device for fast and robust embedding, freezing and longitudinal sectioning of core biopsies (**Figure 1**). The device was designed to ensure functionality and clinical compatibility (**Figure 1a**): This includes the overall sizing and notch at the back (1), which fits into commercially available embedding cassettes and allows their closing for routine labelling (e.g., barcoding) and storage. Furthermore, the U-shaped mold (2) formed by the cover enables precise orientation of a parallel sectioning plane between the longitudinal plane of the biopsy core and the blade before cryo-sectioning. This reduces sample loss due to faster, correct adjustment (**Supplementary Figure S1a**). In particular, a tilted sample would result in sectioning that was not exactly parallel to the longitudinal plane of the biopsy, thus resulting in a smaller measurement area. In addition, the diameter of a biopsy core is very small (usually well below 1 mm), and tissue sections can only be taken from a reasonable depth of the sample (**Supplementary Figure S1b**).

Two different base materials (3), metal and silicone, with different thermal conductive properties were tested with similar results for sample freezing damage (**Supplementary Figure S2**). Both can be sterilized and, in principle, be re-used. Furthermore, both materials are inert and do not react in any way with the sample. Other solutions for aiding biopsy sample preparation include, for instance, filtering paper^34^, which might induce diffusion of liquids out of the sample and therefore change spatial distribution of small molecules. The bridge (4) in the proposed design facilitates transfer of the biopsy core from the needle to the device. This is especially important for biopsy needles including a notch, in which the biopsy resides after excision. Without a punch within the cannula, the biopsy cannot be pushed out of the cannula onto any assisting device. Due to the indentation in which the samples resides, transfer of the biopsy onto a flat surface is impossible, since the needle would mount besides the indentation on the surface and the sample resides in the indentation. For this reason, the bridge with an elevated surface, fits into the indentation of the needle and facilitates release of the biopsy sample onto the device. The cover was made from silicone, which is bendable at negative temperatures (5) and therefore easily removable from the sample. Hence, it allowed for convenient handling while the samples remained frozen in the cryostat. Solid materials like plastics or metal resulted in jamming of the sample in the cover and could not be removed without damage. Learnings made during device development are summarized in **Supplementary Figure S1c – S1f**. The final design consisted of two negative casting molds (made from resin) for the positive silicone parts (**Figure 1b**).

### 3.2. Sample preparation workflow for multimodal, multi omics analysis of needle biopsies

To effectively use the embedding and freezing device, we next developed a sample preparation workflow for consecutive FF and FFPE preparation that includes the following steps (**Figure 2a**): After excision of the biopsy, the core is skimmed off onto the bride of the device’s baseplate (i). By placing the silicon cover on top of the base, a mold is formed around the bridge that can be filled with embedding medium (ii). Embedding also enables handling of non-continuous cores, wherefore no part of the sample is lost. The sample can then be snap-frozen by floating the device in aluminum foil on liquid nitrogen (iii). Before FF biopsy cryo-sectioning, the device is equilibrated in the cryostat, after which the cover is taken off (iv) while the sample remains frozen. Afterwards, the sample is removed from the base and mounted upside-down on a metal chuck (v) for cryo-sectioning. Collected sections can then be used for IRI and MSI (vi) that require FF sample preparation. In addition, one FF tissue section is H&E-stained for reference analysis. A tutorial on the handling of the device for FF preparation up to spectral analyses is provided in the supplementary information. Following cryo-sectioning, the remaining sample is detached from the metal chuck and dissolved in formalin for fixation at RT and for transfer to pathology without sample damage (vii). From here on, routine clinical histopathology analysis including paraffin embedding (viii), sectioning (ix), and subsequent staining is carried out (x; **Figure 2a**).

**Figure 2:**
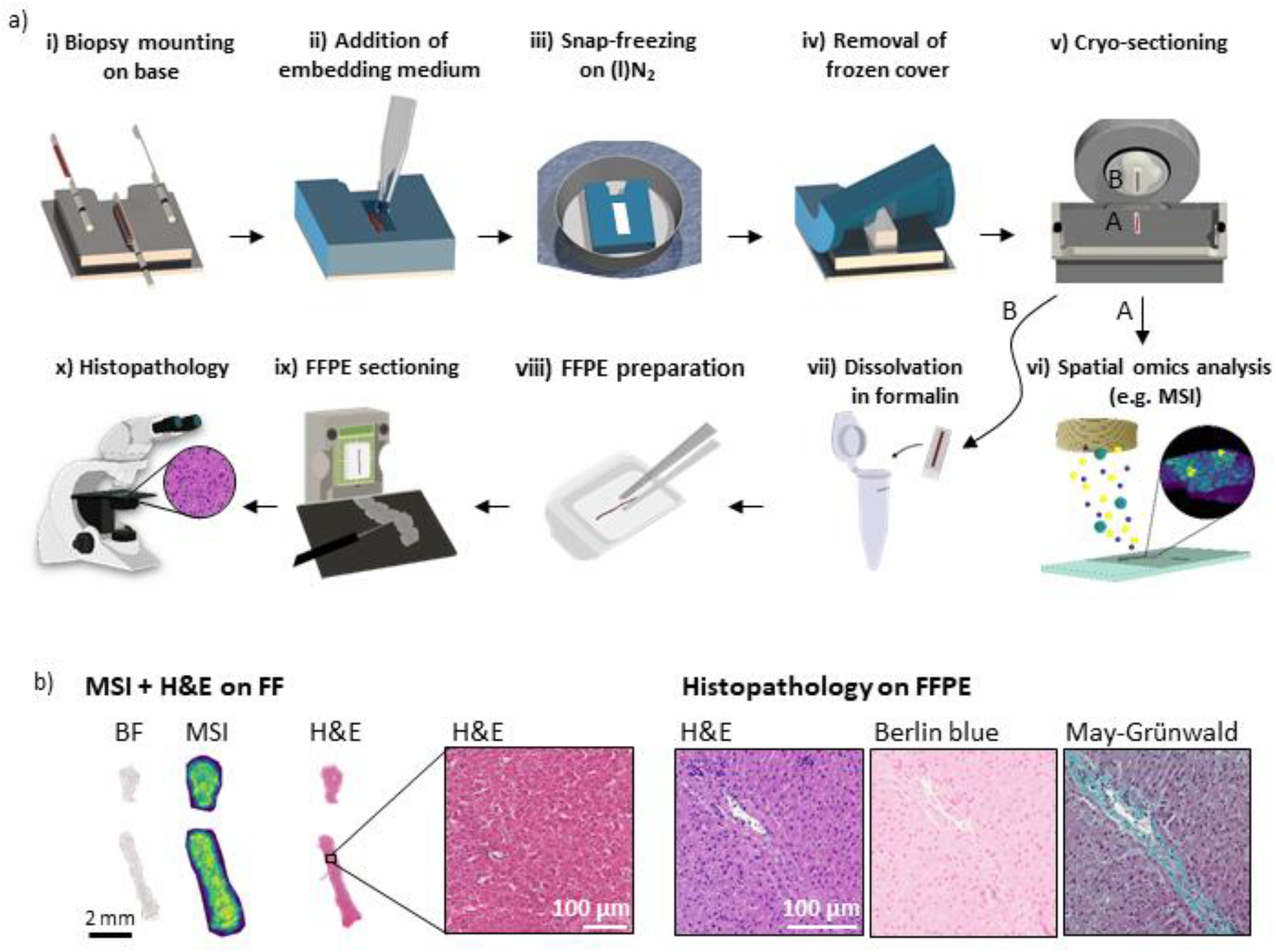
Schematic representation and developmental stage results of workflow for analysis on fresh-frozen tissue sections and subsequent FFPE tissue sections retrieved from a single needle biopsy core. The workflow combines spectral analysis like MALDI MSI and routine clinical histopathology analysis without the need of an additional biopsy. (a) First, the preservation of a freshly excised biopsy sample immediately after needle punching to prevent molecular changes and dislocations in the sample is conducted. This process includes i) transfer of the biopsy from the needle onto the base of the device, ii) embedding in a suitable medium after placing the device cover onto the base, iii) snap-freezing while floating on liquid nitrogen, iv) when frozen removal of the cover (bendable at negative temperatures), v) mounting the sample upside-down on a metal chuck for cryo-sectioning, and mounting sections on suitable slides for the subsequent vi) multimodal spatial omics analyses, e.g., MS- or FT-IRI requiring FF sample preparation. Ensuing the workflow, the same biopsy is preserved afterwards for clinical routine pathology examination, which includes vii) dissolution of the embedding medium and fixation of the biopsy in formalin, viii) embedding in paraffin, ix) FFPE tissue sectioning and mounting on glass slides for subsequent x) histopathological staining and analyses. (b) Exemplary results from bovine liver collected during the developmental stage of the device and according workflow. From left to right: bright field (BF) image, MS imaging (MSI) and H&E staining (all on FF sections) as well as H&E, Berlin Blue, and May-Grünwald stainings performed on FFPE sections retrieved from the same biopsy core following the workflow described in (a).

For multimodal analysis of tissue in spatial omics, several subsequent tissue sections are required. With the presented device and workflow, we retrieved at least eight subsequent sections (of 10 μm thickness) for analyses performed on FF tissue and after subsequent FFPE preparation at least another eight sections (of 2 μm thickness) for analyses on FFPE prepared sections from the core biopsies with a diameter of about 800 μm.

For proof-of-concept work using the described workflow, biopsy needle core samples were initially punched from a bovine liver. In an initial experiment, bright field (BF) images for reference, MSI data, and H&E-stained images were obtained from FF sections prepared from bovine liver biopsies, and H&E, Berlin blue, and May-Grünwald staining were obtained from FFPE prepared sections (**Figure 2b**). This procedure was conducted for two biopsy cores. Multiple additional adjacent FF sections were H&E-stained to visualize tissue morphological quality, similarity of subsequent sections, and to assess registration quality between different modalities for later data fusion (**Supplementary Figures S3 – S5**). Subsequently, these (FF-first) needle cores were FFPE-prepared. Tissue quality and morphology was indistinguishable from a needle core that was directly (i.e. without prior FF procedures) FFPE-processed and thus deemed suitable for accurate pathological annotations (**Supplementary Figure S6**). Using this approach, six out of six biopsies obtained from human samples could be successfully prepared for analysis on FF, as well as FFPE sections.

This demonstrated the robustness of the presented workflow. Moreover, it enabled use of multimodal spatial omics technologies on longitudinally sectioned needle biopsies alongside with clinical routine analysis. This offers the advantage that analysis performed with both preparation techniques are comparable and do not have to be performed on two different biopsy cores, which might be difficult to retrieve from the same patient for ethical reasons, and which might not contain the same tissue due to intra-tumor and general tissue heterogeneity.

### 3.3. Image co-registration for multimodal spectral analyses of needle biopsies

Multimodal analysis often requires different sections for each type of analysis, e.g. MSI, spatial transcriptomics or others, since the techniques are often destructive^12,43^. One common morphological reference for regions of interest (ROIs) are H&E-stained tissue sections with expert pathological annotations. Since sectioning is still mostly a manual process, artefacts such as different deformations of adjacent sections frequently occur, especially in small samples like biopsy cores. To be able to combine results of different sections, one image (the moving image) needs to be computationally transformed to fit the other (fixed image) in a process called image co-registration. Usually microscopic bright field (BF) images used for positional referencing in omics techniques are registered with an adjacent H&E stained ROI reference section. A checkerboard representation of the registration results of two subsequent sections (H&E and BF) provides a quality estimate of registration accuracy (**Figure 3a**). Since non-stained microscopic images do not reveal enough structural detail for the assessment of registration quality, multiple subsequent sections of the bovine liver were H&E-stained. Arteries were annotated as regions of interest (ROIs = landmarks) in each individual section, since these are the most recognizable morphological features in liver tissue, and they usually can be followed across several adjacent sections. These ROI could also be used to determine the overlap between the two registered images and morphological structures, and therefore to assess the quality of the registration result.

**Figure 3:**
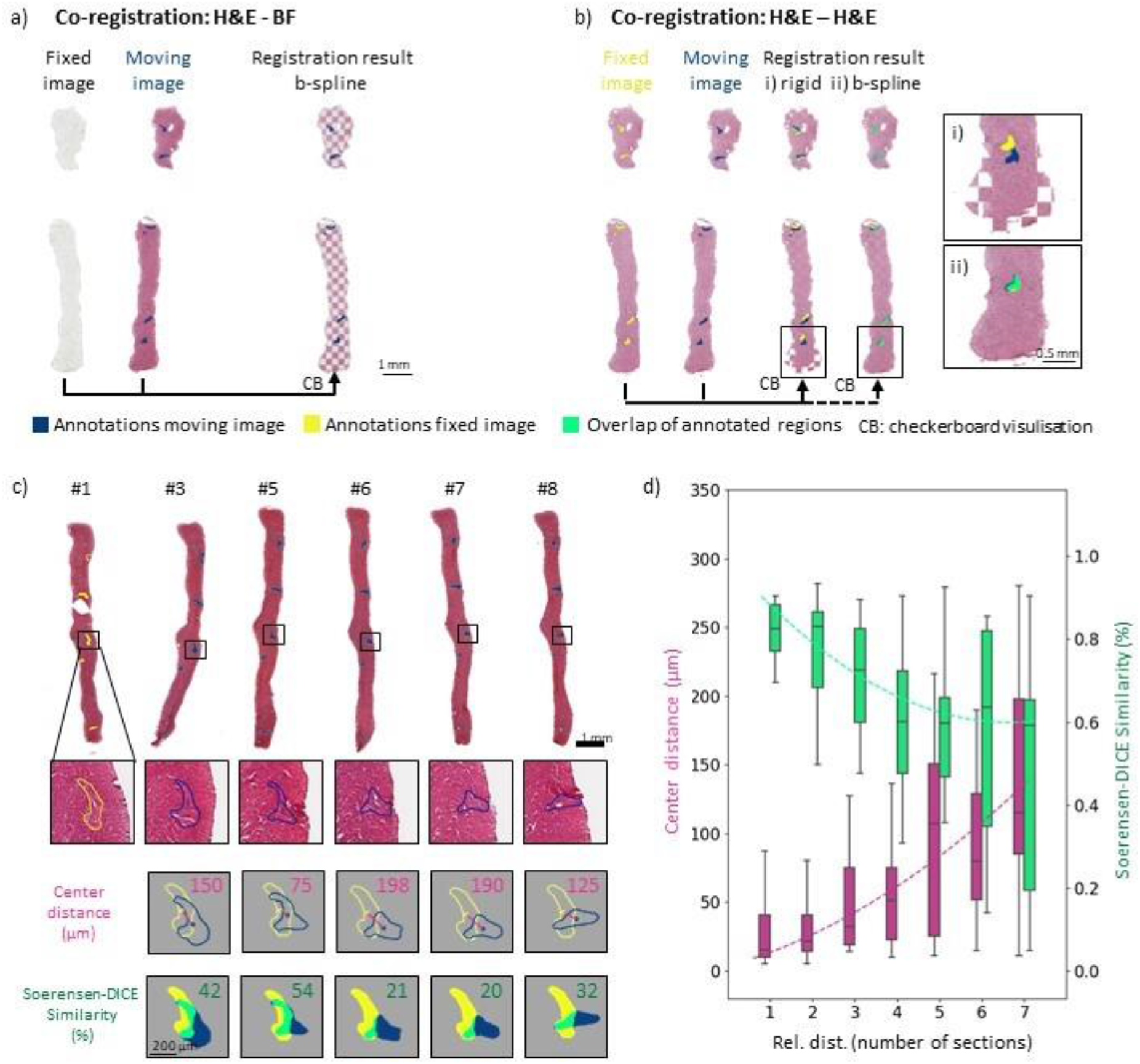
Quality assessment of image co-registration on (FF) bovine liver sections. (**a**) Co-registration of adjacent FF sections, one H&E-stained the other a bright field (BF) image for subsequent spatial omics reference. H&E-stained image was deformed as moving image to fit the fixed BF image with final registration result depicted in checkerboard representation using a non-linear registration method (b-spline). (**b**) Co-registration of two adjacent H&E-stained sections (one fixed, one moving image) with comparison of linear (rigid) and non-linear (b-spline) registration methods, the later allowing for local deformations. Magnifications show the advantage of local deformation possible in b-spline registration, thus enabling a more precise fit of corresponding annotated ROIs (arteries) after the registration. (ROIs: fixed section (yellow), moving section (blue), overlap after co-registration (green)) (**c**) Assessment of registration quality over multiple distances between sections, H&E-stained section numbers (#) (sections #2 and #4 are measured by MSI and were not used for registration quality assessment). (Top) Complete cross-sections of biopsy cores. (middle) Magnification on corresponding ROI (artery) for each section. Note that the shape of the artery changes strongly with progression through the biopsy core. (bottom) Parameter scores to assess registration quality, ROIs gravity-center-distance score and Sorensen-Dice coefficient (SDC) overlaying area score. For each distance the respective section number (#3 – #8, blue ROIs) were moved to fit the fixed section number (#1, yellow ROIs), respectively. Scores are depicted in the top right corners individually. (**d**) Center distance scores (pink) increasing and SDC scores (green) decreasing were calculated for all bovine biopsy samples and their six annotated ROIs and plotted over the relative distance between sections (e.g. distance 1 between sections #5 and #6, distance 2 between sections #5 and #7). Deviations are depicted as box plots (box representing the lower to upper quartile with median representation, outliers lie within the 1.5 x inter-quartile-range (IQR). A second-order polynomial function was fitted to the data to guide-the-eye (depicted as dashed line).

A checkerboard representation was used to visualize the results of two different registration methods, (i) linear (rigid) and (ii) non-linear (b-spline), for two H&E-stained sections (**Figure 3b**). Overlays of biopsy cores after rigid transformation revealed local tissue deformations arising from the sectioning process. Non-overlapping ROI and deviations of the outer shapes of the biopsy cores were observed, as simple rigid registration does not account for local deformations (**Figure 3b i**). Consequently, they could not be matched with this type of transformation method. Accordingly, the ROI of the fixed section image (yellow) and the moving section image (blue) did not overlap (green) well. In contrast, when using non-linear registration (b spline), which allows local corrections including initial linear pre-alignment registration steps, the landmarks and overall shape of the biopsy core were spatially aligned (**Figure 3b ii**; **Supplementary Fig. S6**).

Accuracy of image co-registration depends on the chosen metric. Thus, it is governed by the inherent spatial distribution of histological features present in the sample. Two different metrics for registration quality were calculated for each registration. One was a positional parameter, the median Euclidean distance of the center-of-gravity (center distance) of the annotated features ^41^. It describes the distance between the ROIs after registration. The Euclidian distance, however, neglects a uniform enlargement of the ROIs between two sections, as long as the center-of-gravity remains the same, which describes more precisely the fit of position of the two ROIs after registration. The second metric, the Sorensen-Dice coefficient^41^ (SDC), which evaluates areas rather than positions and which reflects the similarity of two given shapes by their relative spatial distribution, also takes deformations of the ROIs into account.

For multimodal approaches, more than two sections, sometimes adjacent, sometimes with multiple sections in between, need to be registered for comparisons between multiple modalities. For example, tissue sections #3 to #8 were registered (moving images) to section #1 (fixed image; **Figure 3c**). This reflected different numbers of sections that were left out between co-registered sections, e.g., for #1 – #3 a distance of 2, for #1 – #5 a distance of 4 and so on. Sections #2 and #4 were not H&E-stained, but used for MSI instead. An artery, which exhibited morphological changes throughout the biopsy core, was chosen as an exemplarily landmark (**Figure 3c**). The corresponding center distances and SDCs between the landmarks of the moving (#3 – #8) and the fixed image (#1) were calculated. Different types of sectioning artefacts like discontinuous cores, longitudinal stretching of the entire biopsy or bending of the core as well as other examples of annotated arteries and morphological features, which underwent less change throughout eight subsequent sections, are given in the **Supplementary Figure S3 to S5**.

Morphological changes of shape across several adjacent sections and thus center position of the ROIs, deformations of the entire tissue section as well as artefacts introduced during cryo-sectioning significantly impact registration accuracy. For example, some morphological features (like arteries) barely change over eight subsequent sections, whereas others undergo significant positional change (**Supplementary Figure S4b and S7 – S9**). Such effects are not accounted for in linear image co-registration (**Supplementary Figure S6**) and only to some extend even in non-linear transformation. Since the registration used here is intensity-based, the algorithm markedly transforms only those morphological features whose intensity distribution differs from their surroundings (tissue or background). To give one example, best scores for both metrics were achieved for a landmark that presumably represented the cross-section of a vessel (**Supplementary Figure S7 – S9**). High scores for this landmark could likely be attributed to the fact that its shape was nearly constant throughout eight subsequent sections and that contrast between background and surrounding tissue was high. With increasing relative distance of the sections, the center distances increased and SDCs decreased confirming that, not surprisingly, co-registration was more precise for neighboring tissue sections (**Figure 3d**). For small distances between subsequent tissue sections (1 – 3 sections rel. dist.), the median center distance was below 50 μm, i.e. below the size of one MSI pixel (more precisely, its lateral step size). A detailed overview of results for different biopsy samples is presented in **Supplementary Figure S9 and S10**.

### 3.4. Clinical proof-of-concept of FF and FFPE-tissue multimodal spatial omics analysis on the same needle biopsy core

To demonstrate applicability of the proposed device and workflow in a clinical context, samples of two patients were obtained: one diagnosed with hepatocellulary carcinoma (HCC) grade 2 and one with an invasive lobular mamma carcinoma metastasis (MammaMet) residing in liver. Biopsies were not obtained from patients directly, but from freshly resected tissue samples during surgery. Extracted core biopsies (3x HCC, 2x MammaMet) were embedded and frozen using the device and processed via the proposed workflow (**Figure 2a**). For clinical proof of technical feasibility, we aspired to combine the modalities MSI, IRI, and H&E-staining (all on FF tissue sections) and subsequently reference them against histopathological H&E, and IHC staining (all on FFPE sections) conducted on the same needle biopsy core (**Supplementary Figure S9**). Multimodal correlative imaging using MSI, IRI and other modalities has been reported by several laboratories^23,24,26,27,44^. Despite obvious advantages of a single-biopsy-analysis for patients and analytical results, to our knowledge, multimodal spatial omics imaging and classical histopathology have not yet been possible together from the same core needle biopsy processed successively (FF first, then FFPE).

As in initial experiments with bovine liver samples, FF and FFPE tissue quality was suitable for all types of analyses (**Supplementary Figure S2b – c; Supplementary Figure S9**). In some cases, rapture of a biopsy into two parts during subsequent FFPE preparation and sectioning was observed (**Figure 4a vs. c, Figure 5a vs. b**), which, nevertheless, still allowed full analyses and did not affect morphological quality of the tissue, as judged by an expert pathologist. Nevertheless, as the orientation of the biopsy core might slightly change during the transfer from FF to FFPE, precise co-registration between the two sample types is currently challenging. However, due to the small diameter of the biopsy core, it was generally possible to achieve similar cross-sections for FF and FFPE tissue that worked well in comparative bioanalysis. Limitations of the direct comparability between FF and FFPE tissue was only observed for very small ROIs like variances within intra-tumor morphology or differential expression of IHC markers (**Figure 5a and b**). To also enable comparison between such local intra-tumor heterogeneities, the marking of sectioning surfaces during cryo-sectioning would be required for the exact same orientation in subsequent FFPE preparation. This was not attempted in this study.

**Figure 4:**
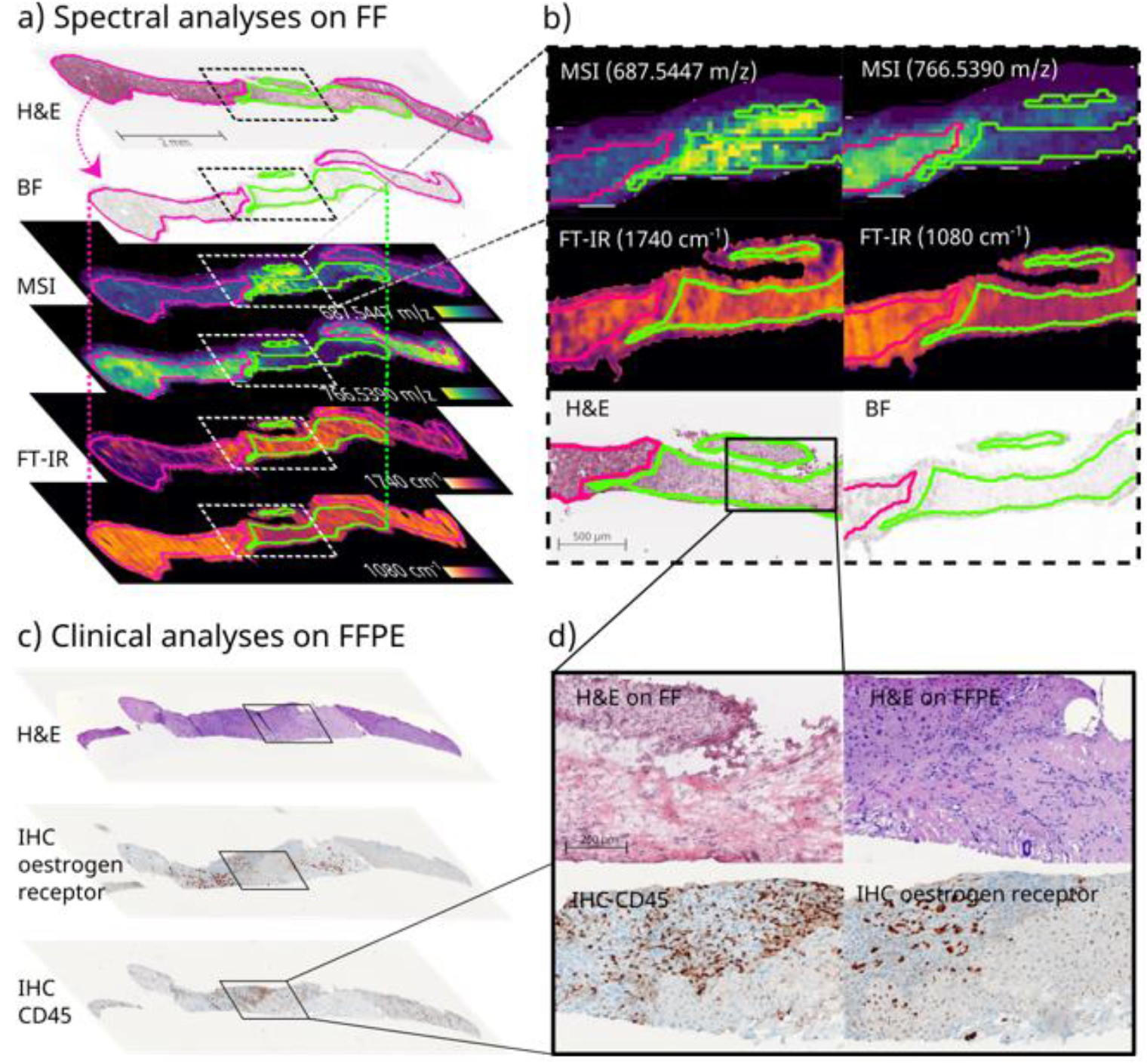
Multimodal imaging on FF and FFPE tissue sections on one biopsy core of a mamma carcinoma metastasis in liver. Following the proposed workflow, FF as well as FFPE sections could be prepared from the same biopsy core for spectral analyses as well as histopathological staining. **(a)** Spectral analysis on FF sections including a H&E stained reference section, bright field (BF) image for registration, MALDI magnetic resonance MSI (spatial resolution 50 μm, *m/z* range 200 – 2000) and FT-IR imaging (spatial resolution 25 μm, spectral range 750 – 4000 cm^-1^). Via co-registration (bent dotted arrows) pathologist annotations (normal liver tissue (pink); liver metastasis of mamma carcinoma (green)) performed on the H&E reference could be transferred (straight dotted lines) onto spectral analysis for discriminant analysis of *m/z* values and wavenumbers. Two examples each of differential distributions between tumorous and non-tumorous regions are depicted including **(b)** magnifications: For MSI, *m/z* 687.545 (± 0.008) displayed higher ion intensity in the tumor region and *m/z* 766.539 (± 0.010) higher ion intensity in the non-tumorous region. For IRI, 1080 cm^-1^ corresponding to P-O-bonds characteristic of nucleic acids and 1760 cm^-1^ corresponding to C-O-double bonds more abundant in lipids were enriched in tumor and non-tumor, respectively. **(c)** Pathological analysis on sections of the same biopsy core after FFPE preparation including H&E, IHC (estrogen receptor and CD45 (lymphocyte common antigen) as a marker of Kupffer star cells) staining with **(d)** magnifications. Additionally, side-by-side H&E staining performed on FF and FFPE sections for comparison of tissue quality, tumor margin, and sectioning plane.

**Figure 5:**
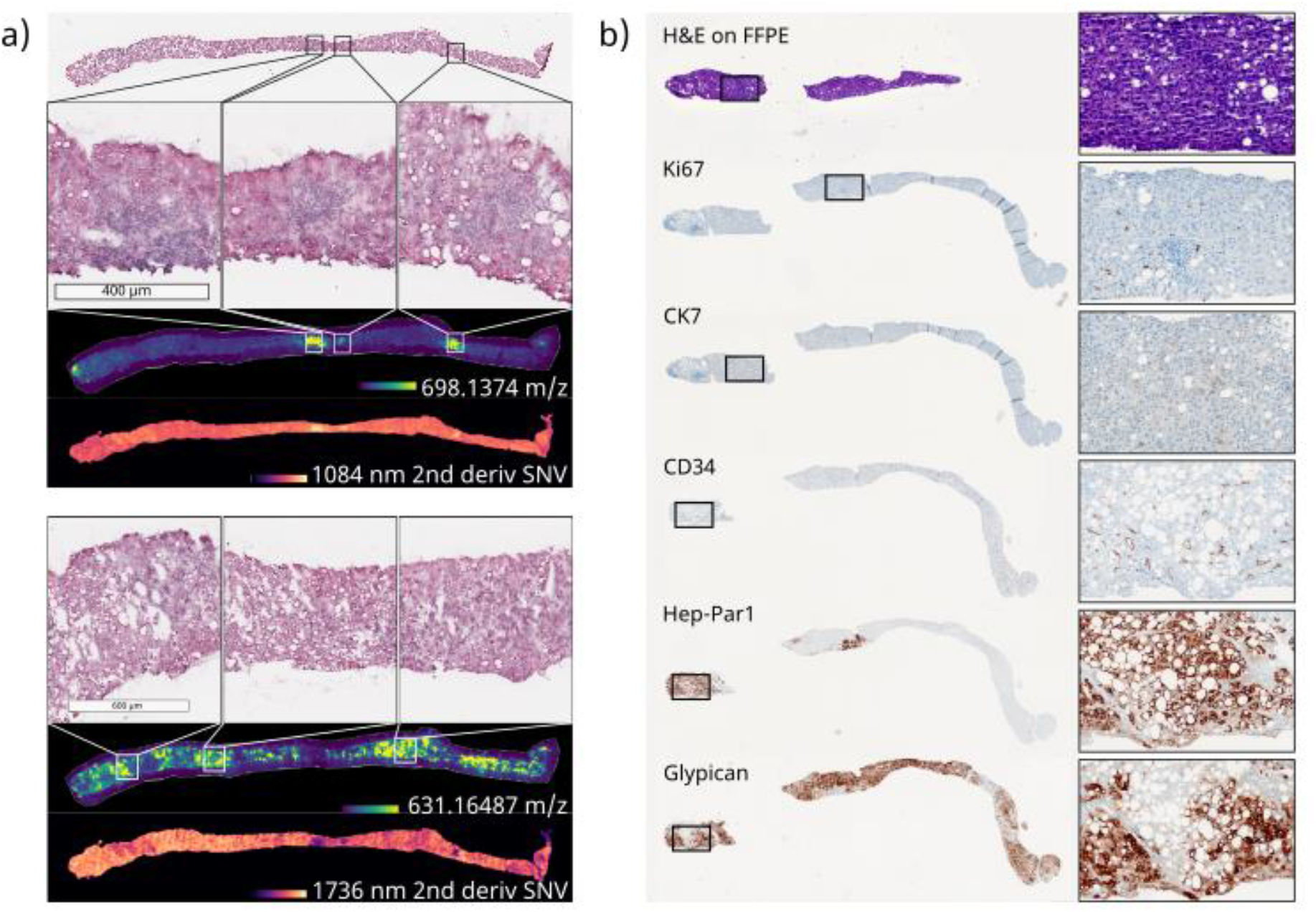
Multimodal imaging on FF and FFPE sections of one biopsy core of a HCC. Following the proposed workflow, FF as well as FFPE sections could be retrieved from the same biopsy core for spectral analyses as well as histopathological staining. **(a)** Spectral analysis conducted on FF sections including H&E stained reference section, bright field (BF) image for registration, MALDI MSI (spatial resolution 50 μm, mass range 200 – 2000 Da, resolution at 400 m/z ^~^135.000, 800 m/z ^~^73.500) and FT-IR imaging (spatial resolution 25 μm, spectral range 750 – 4000 cm^-1^, spectral resolution 4 cm^-1^, pre-processed by second derivative and standard normal variate (SNV)). Pathological annotation suggested the entire biopsy core consisting of HCC tumorous tissue. Nevertheless, different distribution of particular m/z values and molecular classes (locally increased signal intensities) could be observed suggesting intra-tumor heterogeneity. Two examples each of differential distribution are exemplary depicted including magnifications to H&E reference image to correlate morphological variances with the spectral findings. For MSI *m/z* 698.1374 (± 0.008 FWHM), *m/z* 631.1649 (± 0.007 FWHM)) as well as two wavenumbers (1080 cm^-1^, 1736 cm^-1^) indicative of P-O bonds more abundant in nucleic acids and C-O double bonds more abundant in lipids are shown beneath the according MSI images. **(b)** Subsequent FFPE sections are retrieved from the same biopsy core after FFPE preparation and histopathological analysis routinely performed in a clinical context for the precise diagnosis of HCC (CK7, CD34 and glypican3, supported by Ki67 and Hep-Par1 staining) are depicted to emphasize suitable tissue quality for routine analysis on FFPE tissue after beforehand FF preparation.

For this clinical experiment, at least eight sections for both FF and FFPE were prepared, which enabled up to 16 types of analysis on the same biopsy core. In this study, results from MSI, IRI, and H&E staining on FF prepared sections as well as H&E, and several IHC staining on FFPE prepared sections were obtained. H&E staining of FF and FFPE tissue suggested that tumor margins were comparable, despite the expected better morphological integrity of FFPE tissue (**Figure 4d, Figure 5**). Nevertheless, H&E staining on both types of sections were required as references for all analyses done on subsequent sections performed on FF and FFPE tissue. Nuclear estrogen receptor (ER) is diagnostic in this setting, and classification as ER-positive mamma metastasis is supported by additional markers like CK7 (**Figure 4d, Supplementary Fig. S9**).

Generally, the reference is important to describe the content of the presented cross-section, which could include three possible scenarios: only tumor (as observed for the HCC sample, **Figure 5**), a mixture of non-tumorous and tumorous tissue (as observed for the MammaMet sample, **Figure 4**) and only non-tumorous regions (like necrosis or cirrhosis). It should be mentioned that bulk LC-MS/MS Omics approaches that are not spatially resolved are often unaware of the degree of non-tumor tissue in their samples, even though this matter poses a formidable challenge in tumor marker discovery. This can be avoided by using spatially resolved techniques (here, MSI and IRI) and a pathologist-annotated, H&E stained reference sections confirming tissue types.

In line with the primary aim of this study, namely to show the clinical feasibility of the proposed device and workflow, at N=1 no statement can be made about the potential diagnostic significance of observed lipid patterns. With this in mind, lipid annotation in this study (acquired by FT-MRMS in negative ion mode within the *m/z* range of 200 – 2000 with a resolving power of ^~^135,000 at *m/z* 400) used METASPACE (www.metaspace.eu) and a false discovery rate (FDR)-controlled algorithm ^45^. Search against the LipidMaps(2017-12-12) database, resulted in 116 annotations for the mamma metastasis and 4 annotations for the HCC at an FDR of 10% that can be considered level 2 (molecular formulas) annotations according to the Metabolomics Standards Initiative ^46^. Using discriminant receiver-operator characteristic (ROC) analysis in Scils Lab for tumor versus non-tumor ROI, 54 of the annotated 116 *m/z* values in the MammeMet biopsy presented themselves more dominantly in the tumorous region as compared to the non-cancerous tissue. Without this spatial distinction, all 116 *m/z* features would have been attributed to the disease state.

As examples for this differential distribution, two *m/z* features presumably representing two different lipids (*m/z* 687.545 (± 0.008 FWHM), [C38H77N2O6P - H]^-^, 2 candidate molecules with even-numbered fatty acid chains: PE-Cer(d16:1(4E)/20:0) and PE-Cer(d14:1(4E)/22:0)), and *m/z* 766.539 (± 0.010 FWHM/), [C43H78NO8P - H]^-^, 34 candidate molecules), which were predominantly present in healthy and cancerous regions, respectively, are depicted alongside the pathological reference (**Figure 4**). Since the HCC biopsies only presented tumorous tissue, no differential analysis to non-tumorous regions was possible. Therefore, only grouping based on their lipid profile in comparison to a database of tumor entities could reveal relevant molecules for the underlying type of tumor in consistency with other patients having the same diagnosis. Nevertheless, multiple non-annotated *m/z* values showed differential intra-tumor distributions (**Figure 5**). Overall, these results suggested that the proposed device and workflow could be linked with MSI technologies in clinical analysis of minimal invasive samples like biopsies. There would be no need for an extra sample for analysis performed on FF tissue, thus reducing the risk for the patient. These advances in MSI-based cancer research could potentially pave the way for a more comprehensive clinical study with larger patient numbers to achieve more clinically relevant results in form of a complex lipid pattern database accompanying diagnosis or a more profound biological understanding of biological processes in future research.

However, also other spatially resolved techniques like IRI may be beneficial in clinical tissue diagnostics ^26,28,47^. Using annotated ROI (e.g., tumor and non-tumor) as labels, spectral differences can be used to guide clinicians in their disease investigation. In IRI, differential spatial distribution of molecular classes like nucleic acids or lipids is investigated by recording of vibrations of characteristic bonds, like the P-O-vibration in the phosphate backbone of nucleic acids at ^~^1080 cm^-1^ or the higher abundance of C-O double bonds in lipids at ^~^1760 cm^-1^ ^36^. In IRI, these accumulations of molecular classes can be used to some extend as surrogates of e.g. higher proliferation or transcription activity in defined ROI, or of disease states inducing accumulation of lipids as in steatosis. Therefore, differential distribution of molecular classes indicated by the characteristic wavenumbers described above was investigated in this study. These coincided well with the pathological annotations for the MammaMet biopsies between tumorous and non-tumorous regions (**Figure 4**). They also co-localized well with individual *m/z* values suggesting intra-tumor heterogeneity in the HCC sample (**Figure 5**).

In summary, our results show that combined hyperspectral and histopathological analysis of FF and FFPE tissue on the same biopsy core is feasible for the investigation of biomolecular processes in tumors. The ability to combine these modalities even on rather small core biopsies may pave the way to identify tumor-specific markers or signatures. In addition, answering fundamental questions about clinically relevant biological processes may now be possible, if higher samples numbers for each tumor entity were available. Furthermore, using co-registration between hyperspectral and H&E data (in addition to histopathological examination) of the sample should also allow for the implementation of machine learning workflows^48^, e.g., for biomarker discovery^49^ or effective classification models^50^, especially when large-scale studies are performed. Here, the implementation of the presented workflow could be beneficial, since it allows routine clinical collection of biopsy samples, which represents a common source for a higher sample number in the clinic. In addition, it could facilitate early stage diagnosis using molecular, spatially aware digital pathology, perhaps alongside MS approaches in the operating room like intelligent knife technology^51^ or the MasSpec Pen^52^. Overall, multimodal correlative imaging using MSI, IRI and other modalities has been reported by several laboratories^24,44,53^. However, to our knowledge it has not been attempted from the same core needle biopsy after subsequent FF and FFPE preparation yet. The presented approach proved that direct correlation between spectral analyses could be performed using the same biopsy core and therefore correlated with each other to potentially achieve a more comprehensive biological understanding via spatial genomics, transcriptomics, proteomics, lipidomics, and metabolomics describing the entire downstream process ^54^.

## 4. CONCLUSION

In this work, we present a device for sample collection, freezing and sample preparation of core needle biopsy specimen. It enables reliable and robust preparation of tissue sections for multimodal spatial omics analysis on FF tissue in clinical research and complementary routine histopathology on FFPE tissue from the same longitudinally-sectioned biopsy core. Non-rigid co-registration allowed data fusion of various spatial omics modalities from FF tissue for integrative spatially resolved molecular tissue analytics. The proposed approach was successful in an initial clinical feasibility study on five human liver biopsy samples from two different patients with different tumors (namely a mamma metastasis in liver and a HCC). For each sample, at least eight subsequent tissue sections of similar shape and quality could be obtained from each preparation type (FF and FFPE). This approach reduces the risk for patients, since no additional biopsies need to be retrieved for analysis requiring FF tissue. Furthermore, comparability of multimodal measurement results is greatly improved when using the same biopsy core for all analysis, in contrast to using individual biopsies for each analysis. The generation of FFPE tissue from the same core biopsy also permits direct comparison with histopathology, which is the indispensable diagnostic reference for tissue variants, cancer subtypes and disease stages. The device and corresponding workflow is cost-efficient, fast and robust, and therefore may enable standardized implementation in clinical practice. Hence, this standardized workflow appears suitable for clinical routine and clinical research on future early-stage diagnosis, prognosis, and targeted therapy.

## ACKNOWLEGMENTS

The authors thank T. Bausbacher, J. Cordes, A. Geisel, C. Ramallo Guevara and N. Vos for fruitful discussions regarding data evaluation and experimental designs; M. Glaser and L. Schmitt for support with 3D-printing; V. Hartwig, E. Rasbach, B. Fröhlich and T. Soloniewicz for support in the clinic. We thank D. Belharazem-Vitacolonna, S. Hammad and the General Core Equipment unit of the Faculty of Medicine Mannheim, Heidelberg University, for providing central research infrastructure. C. Hopf acknowledges helpful discussions within the COST Action Comulis (Correlated Multimodal Imaging in Life Sciences).

## AUTHOR CONTRIBUTIONS

The study was conceived by M.F.R, S.S., E.B., S.J.D., and C.H. M.F.R. conceived the device, performed sample preparation, measurements, bioinformatics, data analysis, coordinated the work, and wrote the manuscript; S.S. contributed to designing and optimizing the device, performed bioinformatics, data analysis and interpretation, coordinated the work and wrote the manuscript; C.A.W. contributed pathology expertise, annotated tissue sections and provided clinical tissue evaluation; E.B. and N.R. performed surgical resections and provided the human samples as well as clinical information and ethics approval; B.v.M. and M.R. initiated 3D-printing experiments, provided infrared imaging expertise and provided infrastructure. S.J.D. initiated this study and provided infrastructure within the clinical research campus M^2^OLIE. A.M. provided histopathological analysis of tissue samples as well as clinical data and provided infrastructure. C.H. conceived the overall experimental framework, supervised and coordinated the overall work, evaluated results, provided infrastructure and wrote the manuscript.

All authors contributed to, reviewed and approved the manuscript.

## Footnotes

1 https://doi.org/10.1101/2021.10.27.466114

