## supplementary information for "Spatial omics imaging of fresh-frozen tissue and routine FFPE histopathology on a single cancer needle core biopsy: freezing device and multimodal workflow"

VIDEO ON UTILITY OF DEVICE

### Supplementary Methods

**Hematoxilin & eosin (H&E) and immunohistochemistry (IHC) stainings.** For H&E staining of FF samples, sections were mounted on Epredia SuperFrost Plus adhesion slides (Thermo Fisher Scientific GmbH, Braunschweig, Germany), and stained as follows: 2 min hemalaun (Mayer's hemalaun solution, HX14931949, Sigma-Aldrich, Germany), 3 min tap water, 1 min acidic alcohol (350 mL ethanol + 150 mL $H_2O$ + 1.5 mL HCl) (Ethanol absolute EMPLURA®, 818760, Merck KGaA, Germany) (HCL Tritripur, 1.09057.1000, Merck KGaA, Germany), 3 dips in $dH_2O$, 2 min blueing solution (2g $NaHCO_3$ + 20 g $MgSO_4$ in 1L $H_2O$) ($NaHCO_3$, 801K00801129, Merck KGaA, Germany) ($MgSO_4$, 7154.1000, VWR Chemicals, Germany), 3 dips in $dH_2O$, 2 min eosin (Eosin Y-solution 0.5 % aqueous, 1.09844.1000, Merck KGaA, Germany), 3 dips in $dH_2O$, 1 min 80% ethanol, 2 min 96% ethanol, 2 min 100% ethanol, 2 min xylene), and coverslipped with eukitt (03989-100ML, Sigma-Aldrich, Germany) and a glass cover slide (ECN 631-1575, VWR Chemicals, Germany).

H&E and IHC stainings performed on FFPE tissue were conducted to clinical standard procedures. In brief, serial sections (2 µm thick) were taken from FFPE core needle biopsies and stained with H&E, Periodic acid-Schiff (PAS), Masson's trichrome and Prussian blue as well as standard routine immunohistochemistry for diagnostic purposes on a Ventana BenchMark ULTRA, for antigen retrieval a Ventana Ultra CC1 with according retrieval times ($t_{ret}$)., and as a detection system a Ventana OptiView DAB IHC detection kit (Roche Deutschland Holding GmbH, Germany). Antibodies (clones in brackets) were used according to the manufacturers' protocols with corresponding incubation times ($t_{inc}$): CD34 (QBend10, 1:50, $t_{ret}$ of 0 min, $t_{inc}$ of 32 min, Agilent Technologies GmbH, Germany), PanLeu(LCA)/CD45 (2B11+PD 7/26, 1:5000, $t_{ret}$ of 32 min, $t_{inc}$ of 32 min, Agilent Technologies GmbH, Germany), cytokeratin CK 7 (OV-TL12/30, 1:2000, $t_{ret}$ of 48 min, $t_{inc}$ of 32 min, Agilent Technologies GmbH, Germany), HepPar1 (OCH1E5, 1:200, $t_{ret}$ of 32 min, $t_{inc}$ of 32 min, Agilent Technologies GmbH, Germany), glypican-3/GPC3 (1G12, 1:100, $t_{ret}$ of 48 min, $t_{inc}$ of 32 min, Zeta Corporation, USA), estrogen receptor (SP13, 1:200, $t_{ret}$ of 48 min, $t_{inc}$ of 32 min, Invitrogen, Germany), progesterone receptor (pGR636, 1:200, $t_{ret}$ of 32 min, $t_{inc}$ of 32 min, Agilent Technologies GmbH, Germany), ERBB2/HER2 (c-erbB-2 Oncoprotein, 1:800, $t_{ret}$ of 16 min, $t_{inc}$ of 32 min, Agilent Technologies GmbH, Germany), beta-catenin/E-Cadherin (EP700Y, 1:250, $t_{ret}$ of 48 min, $t_{inc}$ of 32 min, Cell Marque, USA) and Ki67 (SP6, 1:2000, $t_{ret}$ of 32 min, $t_{inc}$ of 32 min, Zytomed, Germany). Each IHC staining was conducted using a conjugated horseradish peroxidase (HRP) 2$^{nd}$ antibody and diaminobenzidin (DAB) with a hematoxylin counter staining (32 min total).

### Supplementary Figures

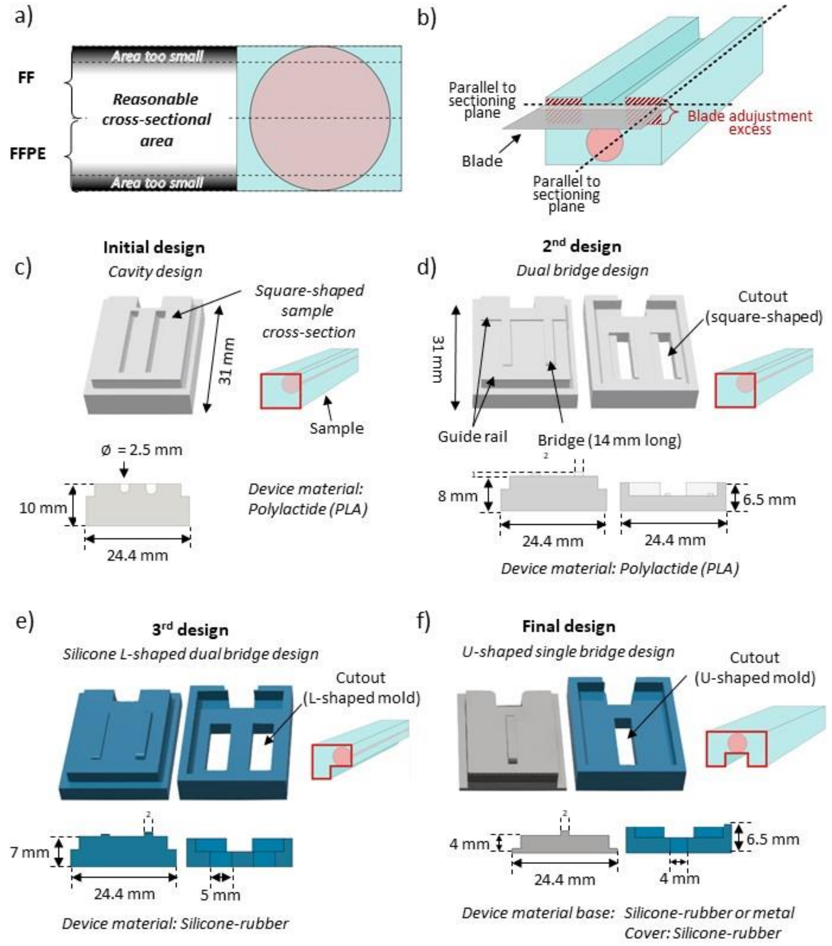

Supplement Figure S1: Evolution of the device design for fine needle biopsy freezing. (a) Sections can only be retrieved from a reasonable depth of the core to obtain enough area for a spatially resolved measurements. (b) The U-shaped cross-section of the embedded sample allowed blade adjustment excess to be visible on both sides of the sample, which allows for precise and faster adjustment of the blade parallel to the sample. Consequently, tissue loss could be reduced. The following developmental steps resulted in the final design: (c) The initial design idea comprised two cavities for embedding biopsy cores and was made out of hard plastic (polylactic acid) by fused-deposition modelling technology using a commercial 3D-printer (Ultimaker2 Extended+, Ultimaker B.V., Utrecht, The Netherlands). With this design, samples could not be removed when frozen. (d) The second design consisted of two parts for better excess to the sample when frozen, and it included bridges to aid skimming off the biopsy cores onto the base plate. Transfer of the sample was improved; however the sample could still not be removed from the top compartment due to expansion when frozen. (e) Therefore, the cover and the base were made from a silicone in the third design, which is bendable at negative temperatures. This allowed easy removal of the sample while remaining frozen. In addition, the biopsy was embedded in a notch of an L-shaped cross-section, which simplified orientation of the sample parallel to the blade. Originally, the device was envisioned with two spaces to hold multiple biopsies at once and save overall storage space and cost. (f) However, since the cover was damaged by mechanical forces (caused by expansion of water-based embedding medium) during freezing, the final design contained only one sample space. With this design, the cover remained intact even after multiple freeze-thaw-cycles and repeated use of the device. Furthermore, the height of the base was reduced to reduce the amount of material between sample and freezing medium and thus reduce freezing damage. For this reason, also two materials of the base plate were tested, silicone and metal, resulting both in sufficient tissue quality after freezing. However, samples could be removed more easily from the silicone base. Hence, this design was used in further experiments. In the final design, also the width of the cutout was enlarged to allow embedding medium to fill the mold on both sides of the sample. This adaptation resulted in a U-shaped cross-section, which allowed further simplified parallel adjustment to the blade (b).

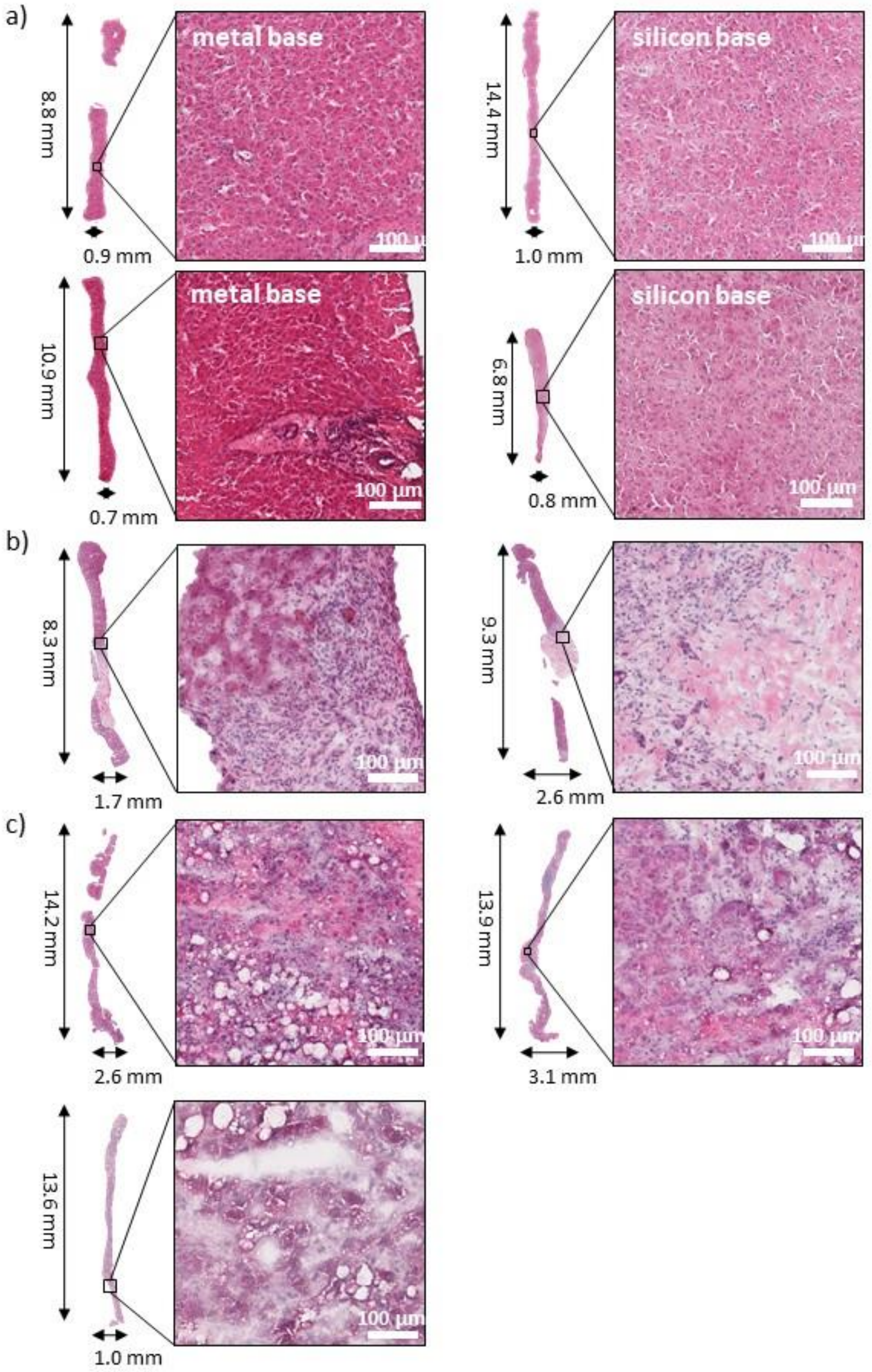

**Supplement Figure S2: Sample overview and histopathological examination of the morphological tissue quality.** (a) Composition of four H&E stained bovine liver biopsy tissue sections used for registration evaluation and examination of the tissue quality. For each base type (silicon or metal), two samples are shown each including a magnified view of a randomly chosen part of the tissue section. Freezing damage between both types of base materials seems to be equal. (b - c) Human biopsy samples included in this study. These include (b) two mamma metastasis in liver, and (c) three hepatocellular carcinoma (HCC) biopsies. Clinical samples were all processed using a silicon base. Morphological quality of the tissue was suitable to assign labels (tumor or non-tumorous tissue) as well as provide a general assumption of the type of cancer.

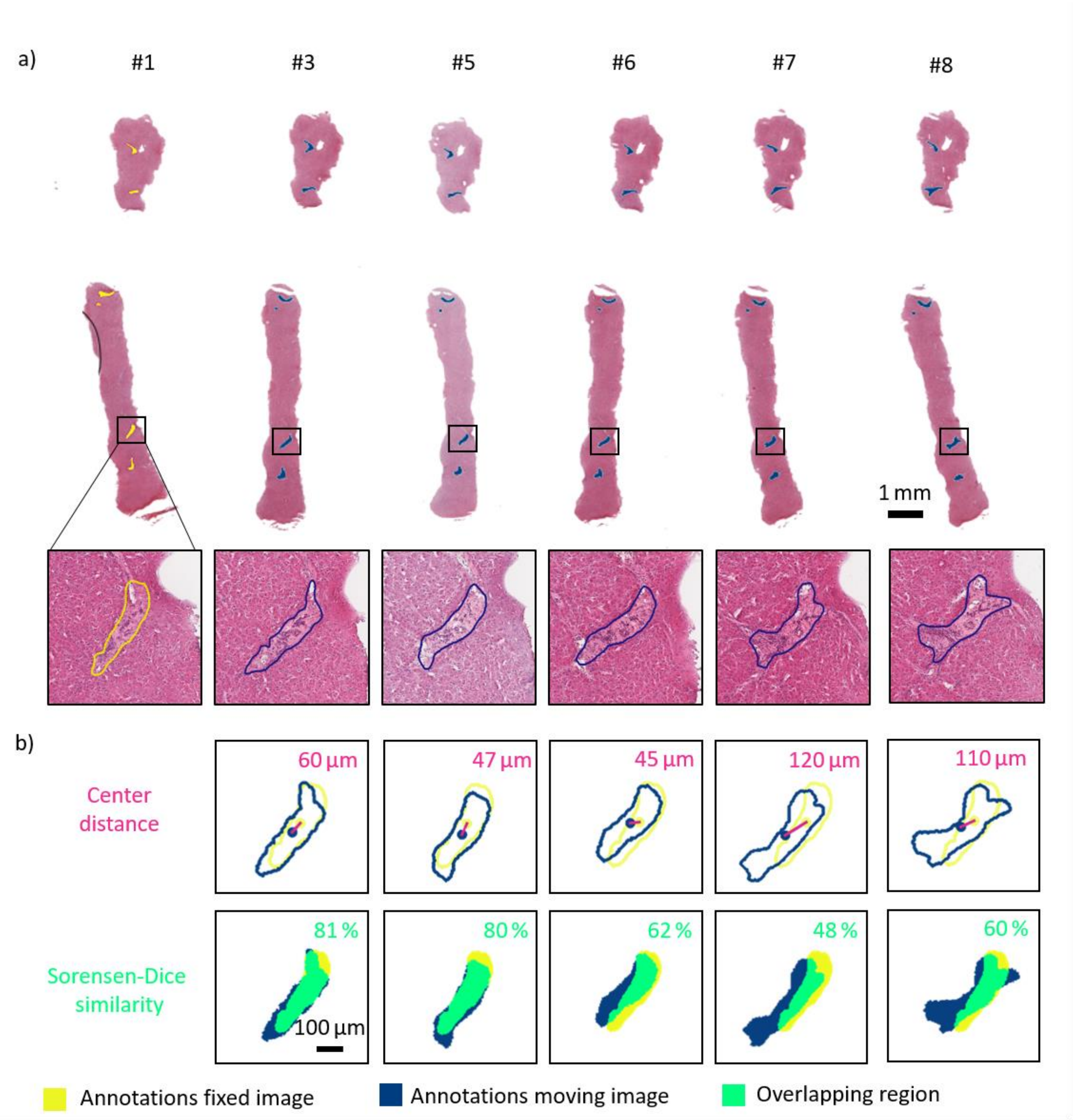

Supplement Figure S3: Co-registered images of subsequent H&E stained bovine liver tissue sections and registration results of a shape-changing annotated ROI. For evaluation of registration, quality eight subsequent sections were H&E stained and several distinctive areas (mostly arteries as these are the most distinctive morphological features in liver) were annotated to determine their fit after registration (non-linear transformation, b-spline). Sections #3 - #8 were deformed (moving image, blue ROIs) to fit section #1 (fixed image, yellow ROIs). (a) Six ROIs spread across the biopsy were annotated as ROIs, which could be observed in each section. Exemplarily, one ROI is magnified to show how shape-changes throughout the sample occur and impact parameters used to determine registration quality. (b) For registration quality assessment, two parameters, the center distance (point-to-point comparison), and the Sorensen-Dice Coefficient (SDC) were used. The center distance is presented as a solid pink line, and the areal overlap of the two regions is highlighted in green. According calculated values of the center distance, as well as the SDC are depicted in the upper right corner for each registration. The peculiarity of this biopsy is that it consists of two parts that are fitted together during registration between the two images.

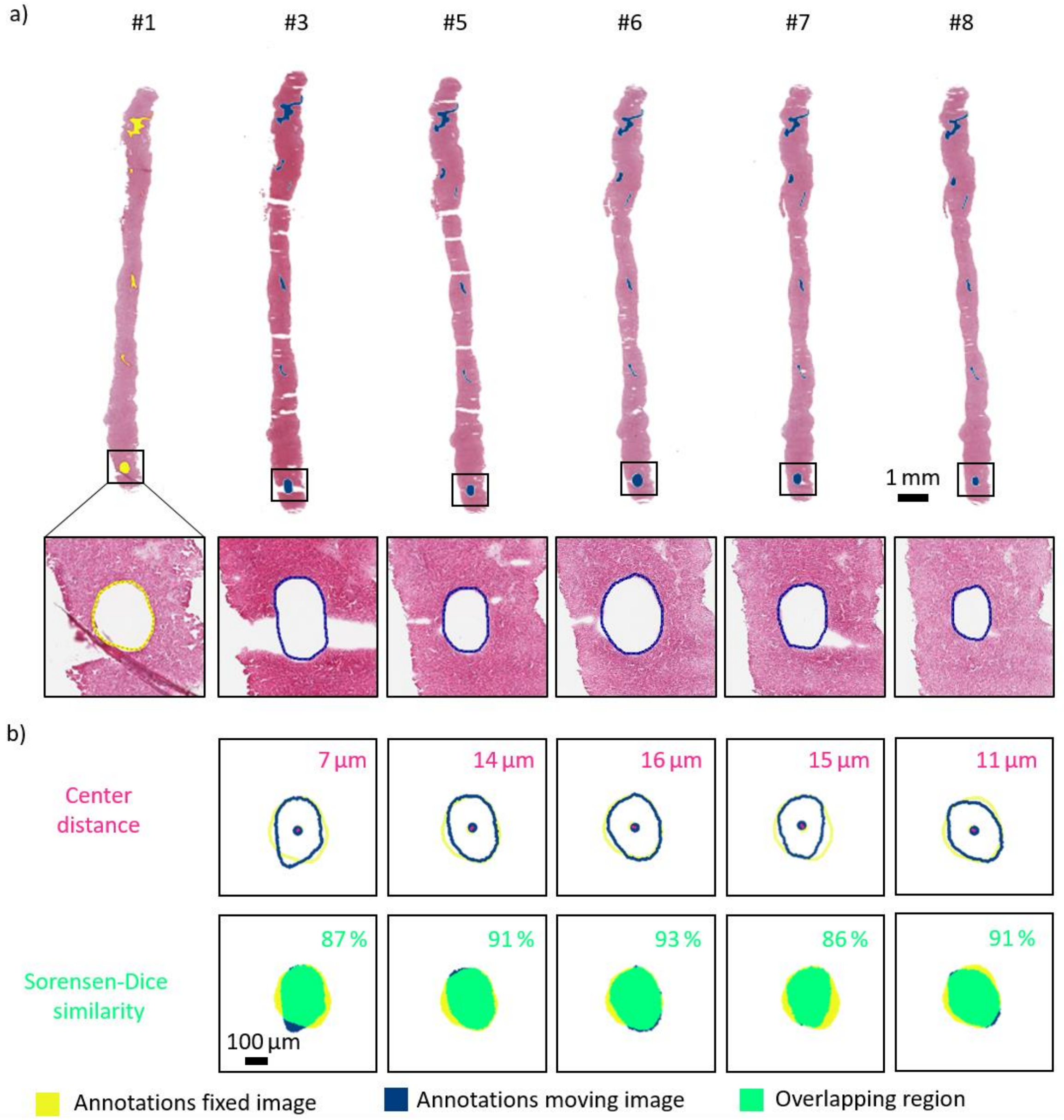

**Supplement Figure S4:** Co-registered images of subsequent H&E stained bovine liver tissue sections and registration results of an annotated ROI with consistent shape. For evaluation of registration, quality eight subsequent sections were H&E stained and several distinctive areas (mostly arteries as these are the most distinctive morphological features in liver) were annotated to determine their fit after registration (non-linear transformation, b-spline). Sections #3 - #8 were deformed (moving image, blue ROIs) to fit section #1 (fixed image, yellow ROIs). (a) Six ROIs spread across the biopsy were annotated as ROIs, which could be observed in each section. Exemplarily, one ROI is magnified to show how shape-changes throughout the sample occur and impact parameters used to determine registration quality. (b) For registration quality assessment, two parameters, the center distance (point-to-point comparison), and the Sorensen-Dice Coefficient (SDC) were used. The center distance is presented as a solid pink line, and the areal overlap of the two regions is highlighted in green. According calculated values of the center distance, as well as the SDC are depicted in the upper right corner for each registration. The peculiarity of this biopsy is that it shows stronger stretching deformations along the entire sample.

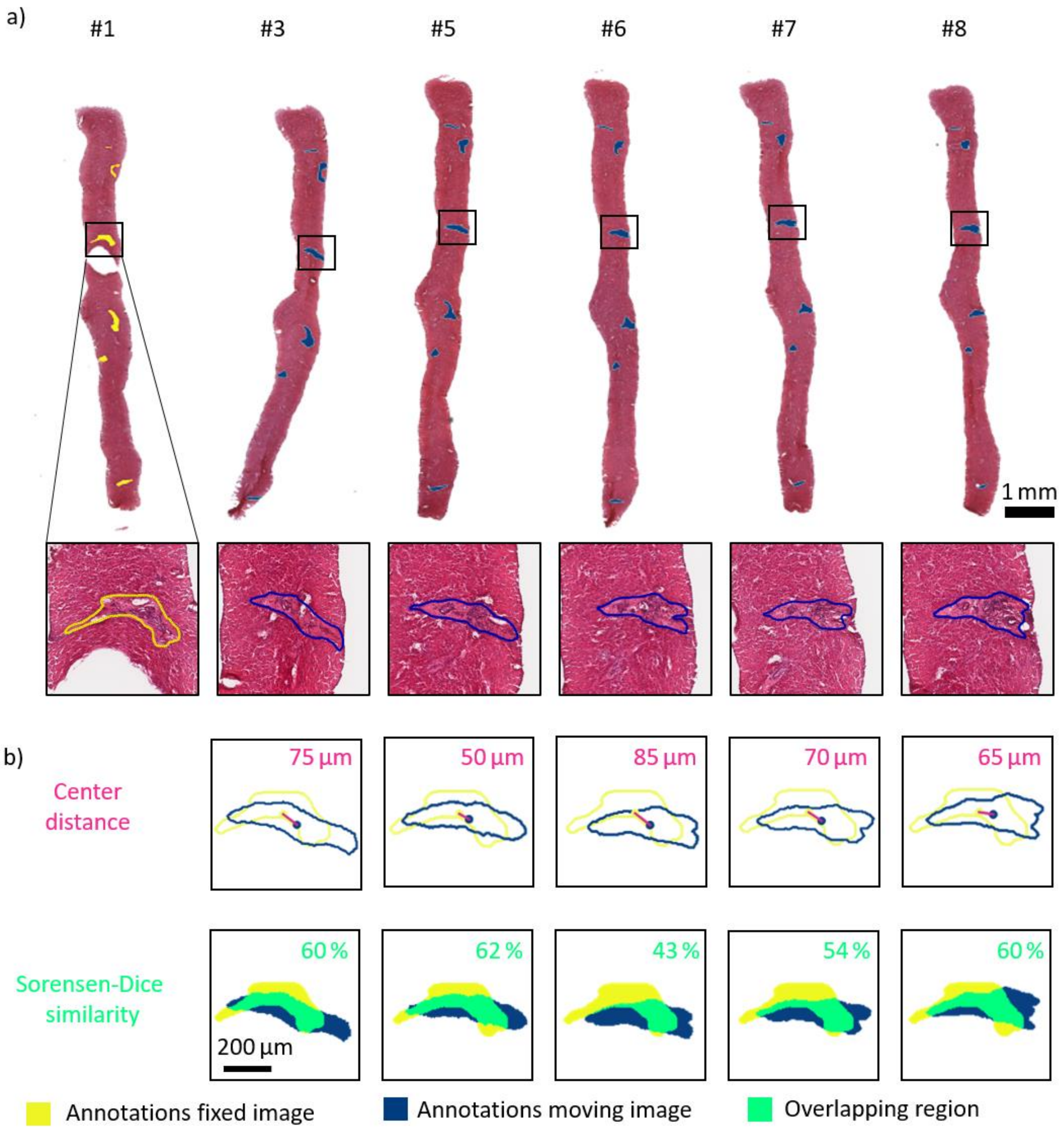

Supplement Figure S5: Co-registered images of subsequent H&E stained bovine liver tissue sections and registration results of a shape-changing annotated ROI. For evaluation of registration, quality eight subsequent sections were H&E stained and several distinctive areas (mostly arteries as these are the most distinctive morphological features in liver) were annotated to determine their fit after registration (non-linear transformation, b-spline). Sections #3 - #8 were deformed (moving image, blue ROIs) to fit section #1 (fixed image, yellow ROIs). (a) Six ROIs spread across the biopsy were annotated as ROIs, which could be observed in each section. Exemplarily, one ROI is magnified to show how shape-changes throughout the sample occur and impact parameters used to determine registration quality. (b) For registration quality assessment, two parameters, the center distance (point-to-point comparison), and the Sorensen-Dice Coefficient (SDC) were used. The center distance is presented as a solid pink line, and the areal overlap of the two regions is highlighted in green. According calculated values of the center distance, as well as the SDC are depicted in the upper right corner for each registration. The peculiarity of this biopsy is that it shows stronger bending deformations along the sample of one section #3 and a missing part of the fixed section #1.

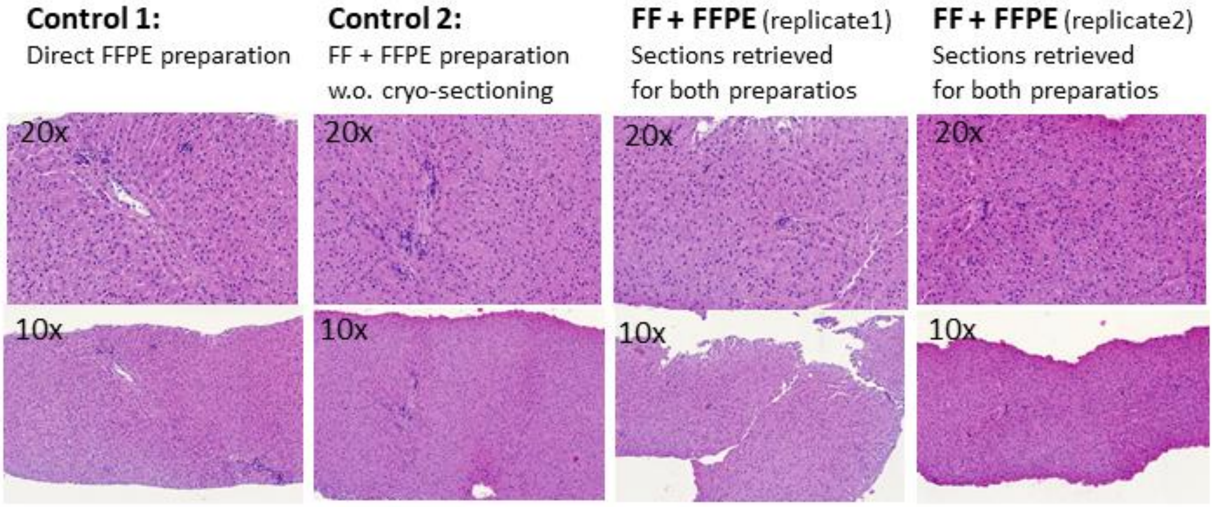

Supplement Figure S6: Tissue quality assessment after FFPE preparation of the initial bovine experiment. Since freezing of tissue samples can induce the reduction of morphological quality a testing experiment, using four liver bovine biopsies was conducted. Two of these biopsies were frozen, sectioned, thawed, and FFPE prepared as described in the proposed workflow (FF + FFPE). Two control biopsies were run alongside from which one was frozen, thawed, and FFPE prepared, but not sectioned to investigate tissue damage, which might occur during the sectioning procedure (control2). The other control biopsy was prepared due to clinical standard procedure and thus not frozen but directly FFPE prepared (control1). This could can be seen as the best possible quality achievable for tissue sections. Tissue quality was observed from sections taken from FFPE prepared tissues and are depicted in two different magnifications (10x and 20x). No difference in morphological quality could be observed between biopsies, which underwent the freeze-thaw-FFPE preparation cycle as compared to the samples directly prepared due to clinical standard procedure. Thus all were deemed suitable for diagnosis by the pathologist.

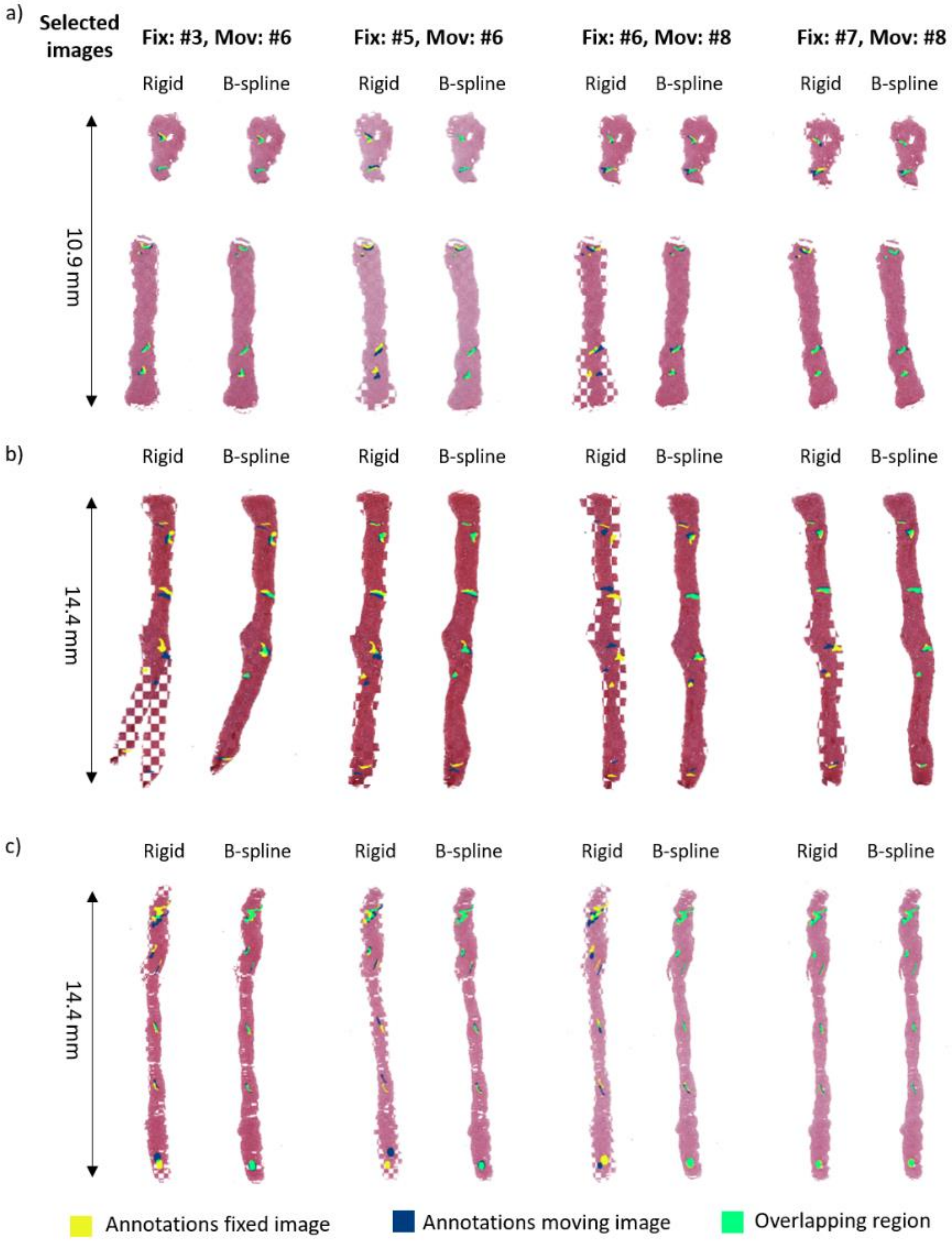

Supplement Figure S7: Checkerboard representation of registration results for linear (rigid) and non-linear (b-spline) transformation. Three bovine biopsies were used to assess registration quality (a – c). For each biopsy four different registrations between varying two sections (section numbers indicated above with #, as well as fixed image and moving (deformed) image assignments). The six annotated ROIs (mostly arteries) of the fixed image were assigned in yellow and according ROIs on the subsequent sections in blue. The overlap between the according ROIs after registration is depicted in green. For a linear (rigid) transformation, which does not allow local transformations of the image, the spatial overlap of the annotated regions is lower as compared to non-linear (b-spline) transformations for all cases as can be observed in the higher abundance of green overlapping areas. Three representations of preparation and sectioning artefacts are depicted here: (a) Non-continuous biopsy core consisting of multiple parts, (b) individual sections that are significantly deformed in one part of the biopsy, and (c) stretching artefact resulting in breaks along the biopsy section.

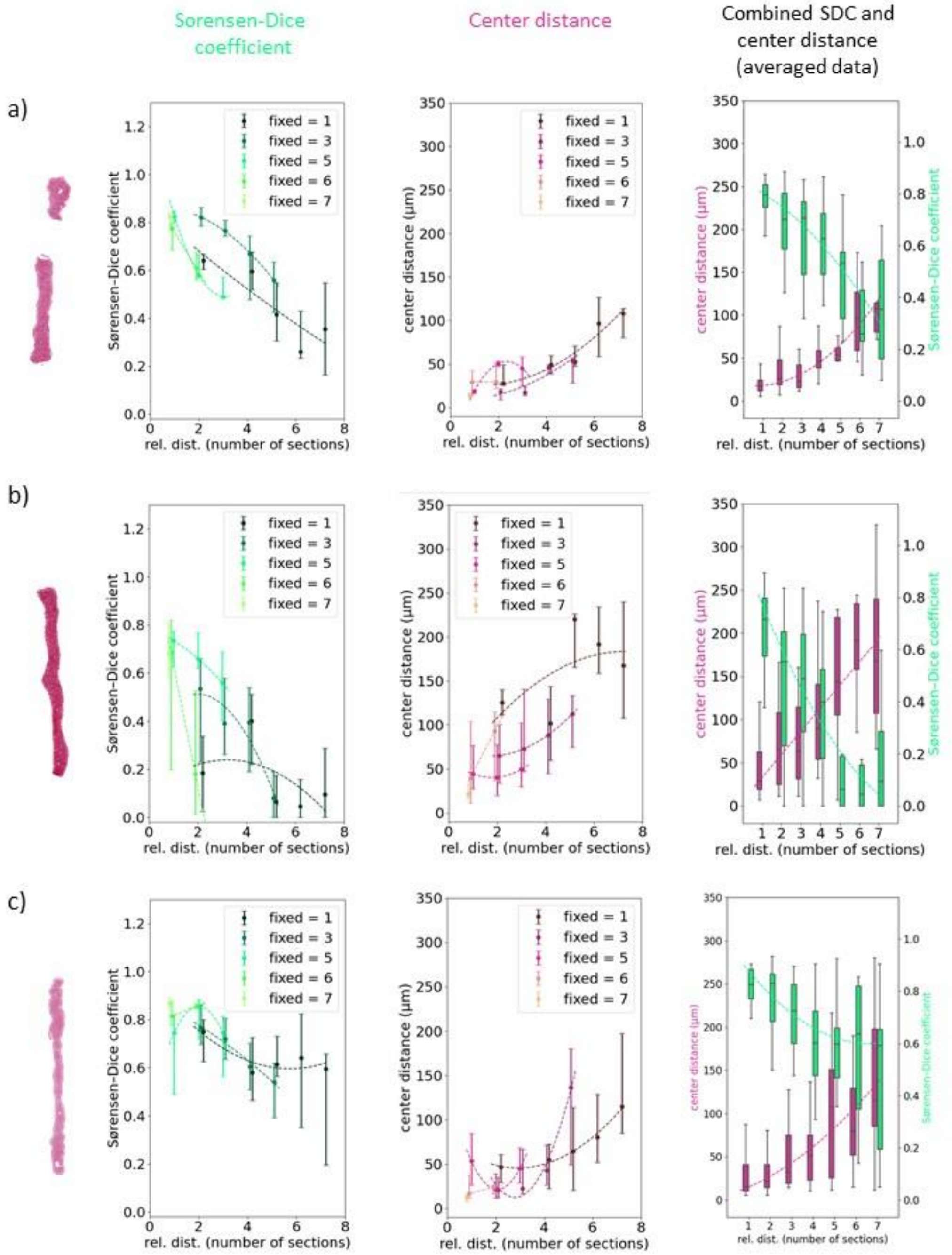

Supplement Figure S8: Registration results for three different bovine liver samples described in SDC and center distance over several serial sections. For each of the three depicted bovine biopsies (a – c) the SDC, as well as the center distance of the six annotated ROIs per section was calculated and their mean with variance (the first and third quantile) is depicted as a function of the distance to their respective co-registered moving image (plots column one and two, respectively). Registration was performed for all possible combinations. Thus, in these plots individual outliers of sections more strongly deformed than others can be explained, which results in the combined plot (plot column three) in a larger variance. Here, averaged results of all registrations within each biopsy sample (a – c) are plotted as a Box and Whiskers plot, with the box representing the lower and upper quartile, as well as the median and whiskers lying within the 1.5x IQR. As expected, in all cases the median values for the center distance increases, whereas the values of the DICE coefficient decrease as a function of the relative distance between tissue sections. In all plots a second-order polynomial function was fit to the data to guide-the-eye.

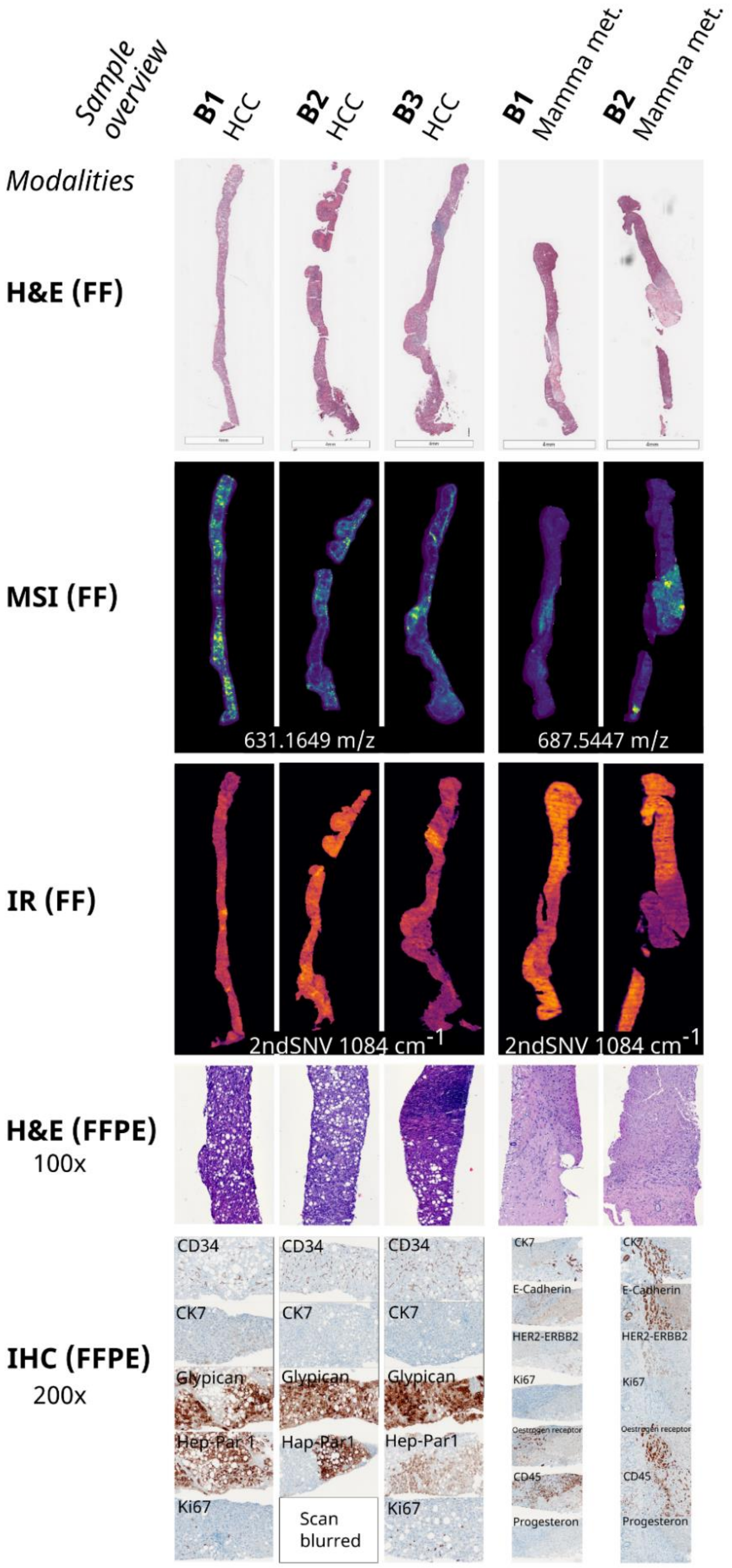

Supplement Figure S9: Overview of human biopsy samples and performed measurements. Images obtained from FF tissue sections include an H&E stained image, a representative MSI image, and a representative FT-IR image. Stainings performed on FFPE tissue sections are an H&E staining (100x magnification) and multiple IHC stainings as indicated (200x magnification) conducted according to pathological routine analysis for the respective diagnosis (HCC biopsies B1 – B3, or MammaMet biopsies B1 – B2).
